# Integrative Microbiome Profiling of Colorectal Cancer Across South Asian and Western Cohorts Using Interpretable Machine Learning

**DOI:** 10.1101/2025.09.23.677677

**Authors:** Mohan Maruthi Sena, Ananya Ganapathy, Viveak Kumar, Ramakrishna Ayloor Seshadri, Narayanan Madaboosi, Mayilvahanan Bose

## Abstract

The incidence of colorectal cancer (CRC) cases has been steadily rising in South Asian countries compared to western countries. Microbiome dysbiosis has been strongly associated with CRC development, with diet and demographic factors playing an important role in shaping gut microbial composition. Since most CRC cases are diagnosed at advanced stages, early detection remains critical for improving patient outcomes. Machine learning (ML) approaches provide a promising strategy to identify predictive microbial biomarkers for early CRC detection. In this study, we have analyzed publicly available 16S rRNA datasets comprising CRC patients and controls from South Asian cohorts (India and Sri Lanka) and compared them with Western cohorts (USA). Our findings revealed a higher relative abundance of *Prevotella* in South Asian cohorts, whereas *Bacteroides* predominated in the USA cohort. Across all three datasets, *Fusobacterium, Escherichia–Shigella*, and *Akkermansia* were consistently elevated in CRC cases. We evaluated multiple ML algorithms, including Decision Tree, Random Forest, AdaBoost, LogitBoost, XGBoost, Support Vector Machine (SVM), and k-Nearest Neighbors (KNN) and proposed stacked ensemble model to differentiate between CRC and controls. Stacked ensemble model has higher accuracy compared to base models. To improve models interpretability and transparency SHAP (SHapley Additive exPlanations) analysis were performed to identify key taxa influencing model predictions. These results underscore the utility of ML-based microbiome analysis for identifying robust CRC-associated microbial signatures, with potential application in developing generalizable early detection tools.

## 1. Introduction

Colorectal cancer (CRC) is the third most commonly diagnosed cancer and the second leading cause of cancer-related deaths worldwide.^1,2^ In 2022, an estimated 1.93 million new CRC cases and 904,000 deaths occurred globally and are projected to reach 4.7 million CRC cases worldwide by 2070.^3,4^ Although historically more prevalent in western countries, incidence is rising in many developing regions, including South Asia. In the recent years, early onset of CRC has been on rise in both men and women across countries.^4–6^ This upward trend may be attributed to demographic shifts, changes in diet, and increasingly sedentary lifestyles. ^7,8^

Diet, demographic factors, genetics, and geographical location are major factors that strongly influence the gut microbiome composition.^9–11^ Recent studies suggests that gut microbiome dysbiosis contributes to the development of CRC progression.^12–17^ To unravel the precise specific contribution of individual bacteria or bacterial toxin studies across diverse populations and contextual factors are essential. However, global microbiome research studies are skewed towards western population, despite rising CRC cases in the developing nations and their representation in global microbiome studies is very low^18^. To address this gap, the International CRC Microbiome Network has been established to promote research in underrepresented regions, including India, Vietnam, Argentina, and Chile.^10,17^ Geographically diverse studies beyond Western cohorts is critical for understanding the global rise of CRC. This will help us to quantify the influence of regional factors on the gut microbiome and lead to accurate prediction of microbial markers for early detection of CRC.

Most CRC cases are diagnosed at intermediate or advanced stages, often after metastasis has occurred.^19–21^ By contrast, early detection can significantly improve patient outcomes, with five-year survival rates up to 90%, underscoring the importance of early detection^22–24^.Therefore, enhancing early diagnostic strategies remains a global priority in CRC research and public health policy.^25^ Recent studies have reported enrichment of specific taxa including bacterial species—such as *Bacteroides fragilis, Prevotella, Escherichia coli, Enterococcus faecalis, Fusobacterium nucleatum*, and *Streptococcus gallolyticus*—in CRC patients compared to controls.^17,26–30 31^ Continued investigation into the gut microbiome’s involvement across diverse population and leveraging on advanced AI techniques could yield valuable biomarkers for early detection and novel therapeutic targets for CRC.

Machine learning (ML) models could assist the identification of potential faecal microbial biomarkers to differentiate between CRC and controls.^12,30,32^ Previous studies have employed machine-learning models to examine the relationship between gut-microbiota composition and CRC, revealing disease-associated shifts in microbial profiles.^17,25,30,32–36^ However, to achieve a deeper and more generalizable understanding of the gut microbiome’s role in CRC, large-scale studies across diverse populations and datasets are essential. Such efforts will help identify both common and population-specific microbial signatures, clarify the contribution of specific microbial species to CRC pathogenesis, and support the development of robust and accurate ML models capable of detecting CRC with high precision.

The current study investigates the microbial diversity associated with CRC in South Asian populations (India & Sri Lanka), faecal profiles from CRC and controls, and comparing against Western datasets (USA) to identify both shared and population-specific microbial signatures. Using relative abundance data, multiple machine learning (ML) models are developed to distinguish CRC cases from controls, guided by optimal feature selection. SHAP (SHapley Additive exPlanations) was applied to the ML models, to identify the most influential microbial features leading to the prediction allowing transparency of the model. These predictive features from each model are stacked into new model to improve the efficiency of the model to identify robust microbial indicators. Taxa that remain consistently high-impact across models and cohorts are proposed as candidate biomarkers, with the longer-term goal of a generalisable early-detection model across diverse populations.

## 2. Methods

### Data Acquisition

We analysed three publicly available 16S rRNA gene (V4 region) amplicon datasets from India (ENA accession id: PRJEB53415; 43 CRC and 46 controls (healthy volunteers)), Sri Lanka (ENA accession id: ERP161747; 24 CRC and 50 controls), and the USA (ENA accession: PRJNA290926; 120 CRC and 171 controls)^17,27,29^ These datasets were selected on the basis that they have employed comparable experimental protocols for DNA extraction from faecal samples.

### Data processing

Adapters from the downloaded reads were removed using cutadapt.^37^ The reads devoid of adapters were verified for quality control using Fastqc.^31^ To ensure high-quality data for downstream analysis, reads were trimmed to a uniform length of 220 bp. Thus, obtained high quality trimmed reads were further processed using QIIME2 (version 2024.5).^38^ Denoising was then performed using DADA2 (version 2024.5) within the QIIME2 pipeline to generate representative amplicon sequence variants (ASVs) for downstream analyses.^39^

Taxanomic classification was performed using SILVA database (version 132)^40^ using BLAST+^41^ implemented within QIIME2 (version: 2024.5) *q2 feature classifier* plugin.^42^ The sequences were rarified to the lowest depth sample 43000 (QC passing reads, India), 2965 (Sri Lanka), 1942 (USA) before diversity analyses. Alpha diversity was assessed using Shannon index,^43^ while beta diversity was evaluated using Bray-Curtis dissimilarities.^44^ Associations between Bray–Curtis distances and sample metadata were tested with Permutational Multivariate Analysis of Variance (PERMANOVA) via the *adonis* function.^45^ Principal Coordinate Analysis (PCoA) was used to visualize beta-diversity patterns.Taxa tables, diversity metrics were exported from QIIME2 for subsequent analyses and visualized using R (version: 4.3.0). Machine learning models, including Random Forest (RF),^46^ XGBoost,^47^ AdaBoost,^48^ LogitBoost^49^, and Decision Tree^50^, SVM,^51^ KNN^52^ were constructed using the scikit-learn package in Python (version: 3.12.2) and their performance was validated using the pROC.^53^ Linear discriminant analysis effect size (LEfSe) was employed to identify specific taxa significantly associated with different metadata categories.^54^

## 3. Results

### Beta Diversity

The principal coordinate analysis (PCoA) plots for the three datasets are shown in Figure 1. The clustering pattern revealed stronger separation by geographic origin than by cancer status, consistent with previous studies.^17^ The plot is dominated by the western samples as the size of the dataset is more, the western samples are located on the right of the plot, South Asian samples to the left. Adonis PERMANOVA analysis indicated variation of 42.5% with country of origin and cancer status 1.8 % (p < 0.001).

**Figure 1:**
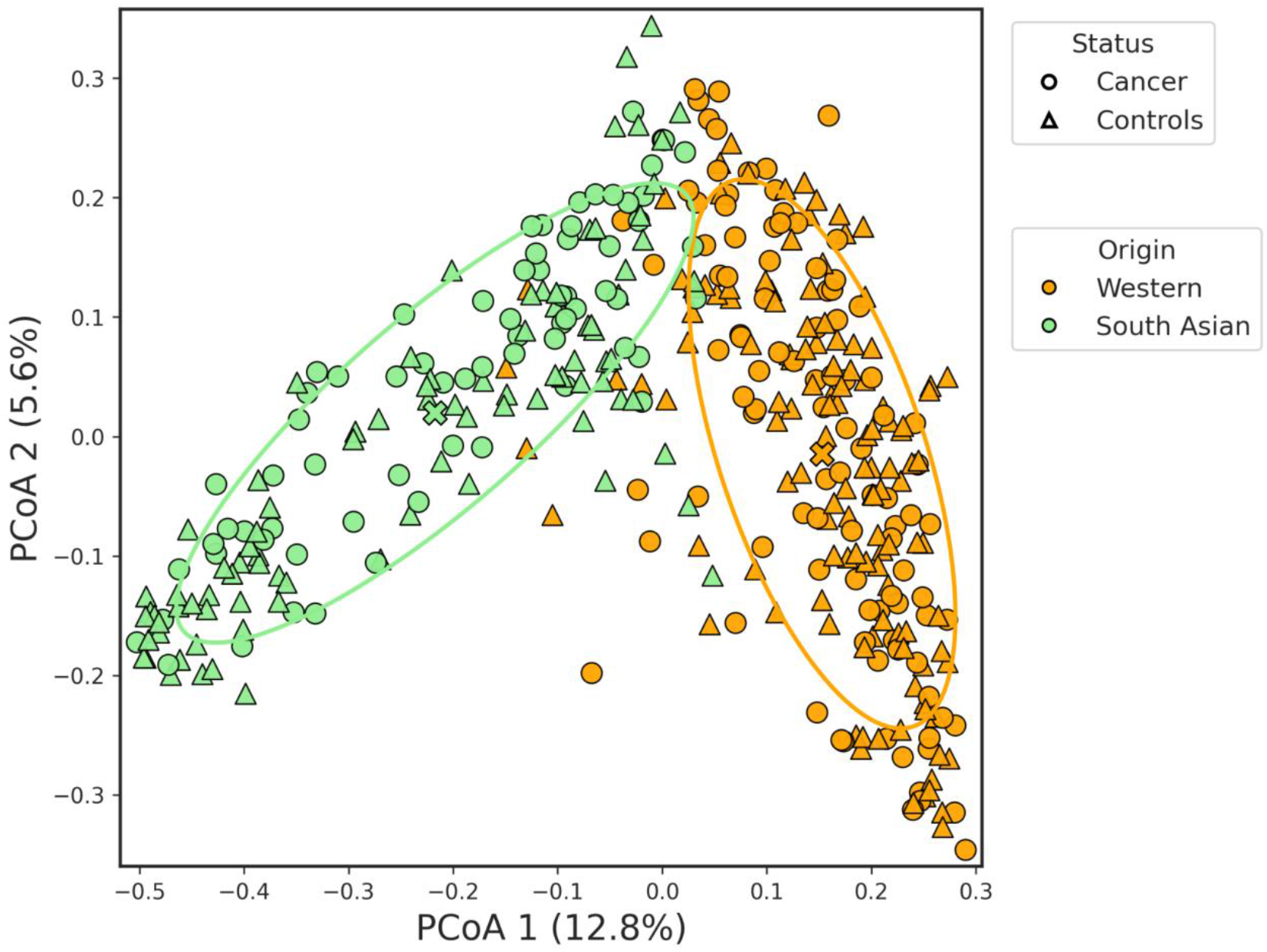
Principal coordinate plot of Bray-Curtis distances for the datasets considered in this study. 95% confidence intervals for sample groupings are displayed as ellipses.

The alpha diversity across all the datasets considered in this study is shown in Figure 2. Western samples (USA) have higher diversity compared to South Asian (India+Sri Lanka) samples. Within each dataset, cancer samples had higher alpha diversity than controls, consistent with earlier reported studies. Although the sample sizes vary across different cohorts, the trend was confirmed statistically (Mann-Whitney p=0.0005).^17,26^ Supplementary Figure S2 shows the alpha diversity of samples from India and Sri Lanka, revealing that cancer samples have a higher diversity than samples.

**Figure 2:**
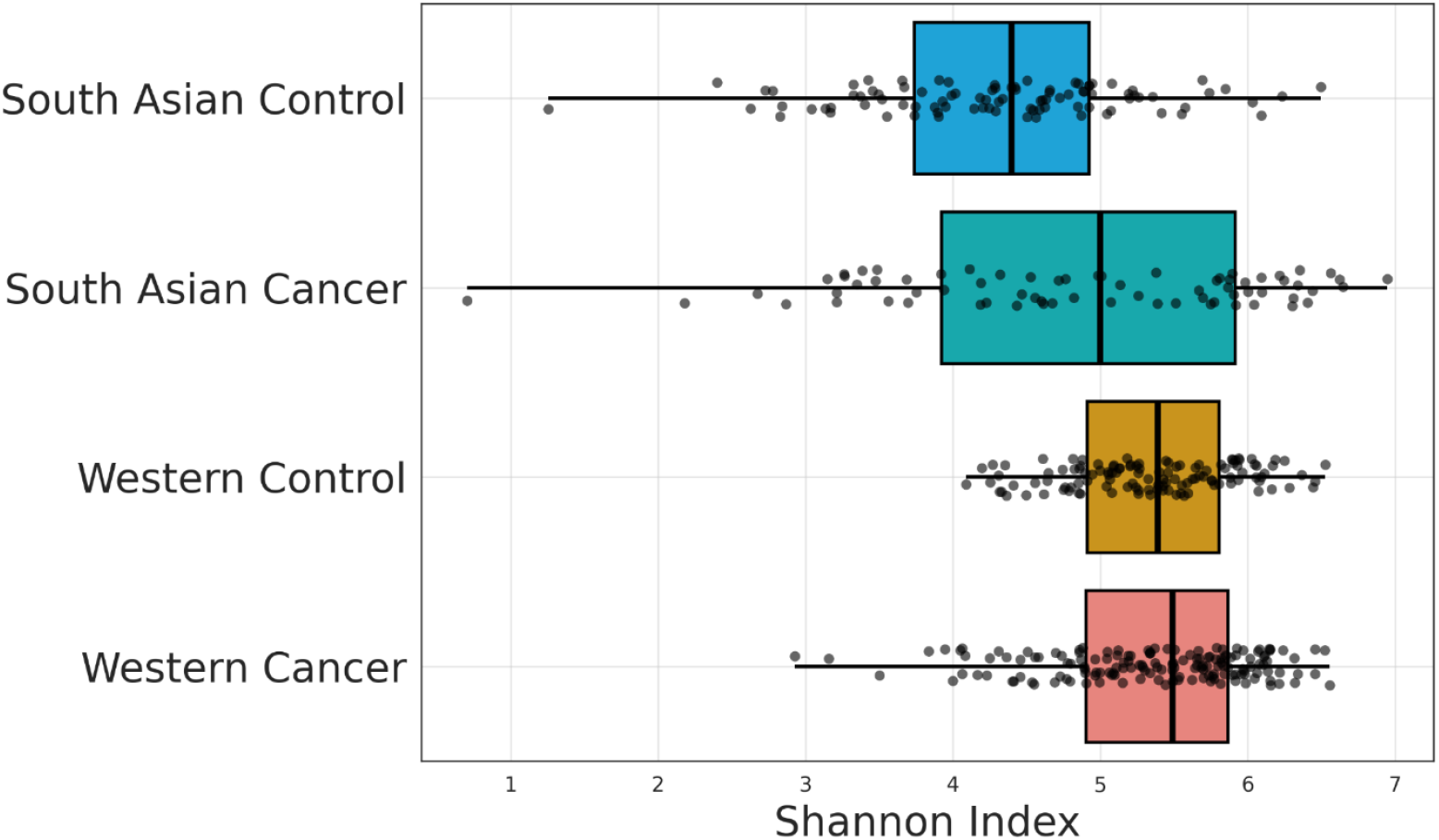
Shannon index alpha diversity for South Asia and Western cohorts.

### Taxa Abundance

The top 15 dominant microbial groups across the datasets considered revealing a distinct separation based on geographical location consistent with the PCoA results (Figure 3). South Asian populations exhibit a higher abundance of *Prevotella*, while Western populations show a greater prevalence of *Bacteroides*. Notably, the Sri Lankan cohort, unlike the Indian cohort, displays a higher abundance of *Bacteroides (Figure S2)*, underscoring the influence of regional variation on gut microbiome composition.

**Figure 3:**
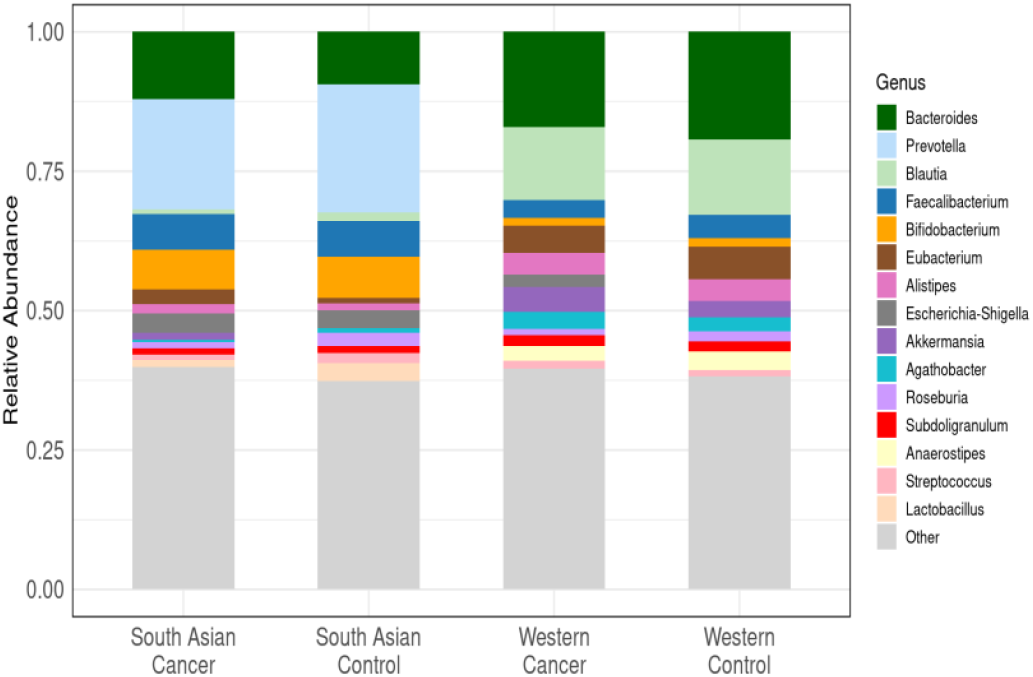
Taxa abundance showing top 15 genera across all the datasets considered.

Among the top genera, *Escherichia/Shigella* are more prevalent in cancer cohorts. To identify microbial differences between cancer patients and controls, taxa from all countries analyzed using LEfSe (Figure S3-S5).^55^ This analysis, which extended beyond the most common genera, identified *Akkermansia, Fusobacterium*, and *Ruminococcus* as prominent in CRC-associated fecal flora. A complete list of Lefse analysis for the datasets considered can be found in Figure S3-S5.

## 4. Dataset Preparation for Machine Learning

The input dataset for training the machine learning (ML) model consists of genus-level microbial abundance tables derived from 16S rRNA V4 region sequencing of samples from South Asian (India, Sri Lanka) and Western countries (USA). Preprocessing involves removal of unclassified taxa (‘Other’) from the genus tables and normalisation of genus-level abundances across all samples. To enrich the feature space for classification, we integrated available metadata—age, BMI, smoking status, and gender (Western and Indian datasets) were incorporated into the dataset. The resulting pre-processed data were subsequently used to train a suite of regression and classification models.

### 4.1 Machine learning classifier

#### 4.1.1 Random forest

The Random Forest (RF) algorithm is an ensemble learning technique used for both classification and regression.^56^ It builds multiple decision trees during training, forming a “forest.” For classification tasks, the final output is based on the majority vote from all trees, while for regression, the result is the average of their predictions. Compared with a single decision tree, RF typically reduces overfitting, handles high-dimensional feature spaces and nonlinear interactions, and provides feature-importance estimates.

In this study, we optimised a CRC classifier using k-fold cross-validation and a grid search over RF hyperparameters: max_depth ∈ {None, 3, 5} and n_estimators ∈ {50, 100, 150, 200, 250}. The model was implemented in Python 3.9.21 using scikit-learn 1.6.1.^57^

#### 4.1.2 XG (eXtreme Gradient Boosting) Boost

XGBoost (Extreme Gradient Boosting) is an ensemble learning method that constructs decision trees sequentially, where each new tree aims to correct the errors made by the previous trees.^58^ It utilizes gradient descent to minimize a specified loss function and incorporates several key enhancements over traditional gradient boosting algorithms, including regularization, parallel processing, and efficient tree pruning.

In this study, the following hyperparameters were tuned for optimal performance:

a. max_depth ∈ {3, 5}
b. n_estimators ∈ {50, 100, 150, 200, 250}
c. colsample_bytree ∈ {0.2, 0.4, 0.6, 0.8}

The XGBoost algorithm was also implemented using Python (version 3.9.21) and the XGBoost library (version 2.1.4).

#### 4.1.3 AdaBoost (Adaptive Boosting)

AdaBoost (Adaptive Boosting) is an ensemble learning technique that combines multiple weak learners to form a strong predictive model. It works by iteratively improving the model’s performance, placing greater emphasis on data points that were misclassified in previous iterations. Initially, equal weights are assigned to all training samples; these weights are then adjusted after each iteration to focus more on the incorrectly classified instances, thereby guiding the learning process in subsequent rounds.

In this study, the following hyperparameters were tuned to achieve optimal performance:

- max_depth ∈ {3, 5}
- max_iter ∈ {500, 1000}
- n_estimators ∈ {50, 100, 150, 200, 250}
- learning_rate ∈ {0.2, 0.4, 0.6, 0.8, 1.0}

#### 4.1.4 LogitBoost

LogitBoost is a statistical variant of the AdaBoost algorithm, which differs by directly minimizing the logistic loss function.^59^ While the core iterative boosting framework remains similar to AdaBoost, LogitBoost integrates a probabilistic approach more suited for classification tasks by modeling the log-odds of the target variable.

In this study, the following hyperparameters were optimized for best performance:

- max_depth ∈ {3, 5}
- max_iter ∈ {500, 1000}
- solver ∈ {liblinear}

#### 4.1.5 Decision Tree

Decision Tree is a supervised machine learning algorithm that recursively partitions the dataset into homogeneous subsets based on feature values.^60^ At each node, the algorithm selects the optimal feature and threshold that best separates the data according to a chosen splitting criterion (e.g., Gini impurity or entropy). This hierarchical structure makes decision trees highly interpretable and effective for both classification and regression tasks.

In this study, the following hyperparameters were tuned to optimize model performance:

- max_depth ∈ {3, 5}
- splitter = ‘best’ (selects the feature that yields the highest information gain)
- max_iter ∈ {500, 1000}

#### 4.1.6 Support Vector Model (SVM)

The Support vector machine is a supervised machine learning algorithm widely used for binary classification tasks. It segregates the data by identifying a hyperplane that minimizes the loss function. The Data points lying close to the hyperplane act as supporting vectors that contribute to define the boundary. For the classification of CRC/normal, we have used cross-validation along with the tuning of following parameters for optimization,

- C ϵ {1, 5, 10, 20},
- Gamma ϵ {0.001, 0.01, 1}

The SVM algorithm is implemented using the Python (version 3.9.21) package scikit-learn (version 1.3.0).^61^

#### 4.1.7 K-nearest neighbor (KNN)

KNN is a supervised machine learning method used to classify data based on the nearest neighbours. As a non-parametric and instance-based method, it relies on the principle of similarity to classify a new data point based on the majority class of its ‘k’ nearest neighbors in the feature space. The choice of ‘k’, the number of neighbours, is a critical hyperparameter that significantly impacts model performance. The values k∈{5,10,15,20} were likely evaluated to find the optimal balance between bias and variance for the given problem.

### 4.2 SHAP Algorithm

SHAP (Shapley Additive exPlanations) provides interpretability and transparency to machine learning models that are often perceived as “black boxes.” This method is based on game theory, SHAP explains individual predictions by assigning each feature an importance value—known as the SHAP value— representing its contribution to the model’s output.^62,63^ This method decomposes a prediction into additive contributions from each feature, ensuring that the sum of these contributions equals the difference between the actual prediction and the average prediction across all samples.

In this study, we employed the SHAP framework to identify and quantify the influence of individual features in distinguishing between controls and CRC samples. The model F requires retraining on all subsets F of the complete set S of features (F ⊆ S). The SHAP value for the j^th^ feature of the instance x is determined by aggregating it across all possible subsets (Equation 1)

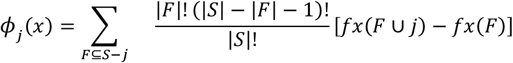

Where |F|! represents the permutations of features in the subset F, (|S|-|F|-1)! The permutation of features in the subset S – (F ⊆ {j}) and |S|! is the total number of feature permutations.

SHAP value computations were carried out using the Python package SHAP (version 0.48.0) under Python 3.9.21. For tree-based models including Random Forest (RF), XGBoost, and Decision Tree, we utilized the TreeExplainer with the feature_perturbation parameter set to “interventional” to capture robust feature impact estimates. For AdaBoost and LogitBoost models, the KernelExplainer was applied due to their non-tree-based structure. This strategy was particularly useful in evaluating the impact of highly correlated predictors and mitigating potential misinterpretations.^64^

### 4.3 Key Features for Cancer Status Prediction Using ML models

In this study, we have employed various ML models including Random forest, Logitboost, Adaboost, XGboost, Decision tree, Support Vector Machine (SVM) and K-Nearest Neighbours (KNN) to discriminate between CRC and controls using features selection. The top 20 discriminative features for the identified respective algorithm shown in the Figure S5-S17. Two complementary strategies were applied to build the model,

#### i) Baseline Models using all features

The above mentioned models were trained using the genus level tax abundance from South Asian (India+ Sri Lanka), and western cohorts (USA) without applying feature selection.^34,65^

#### ii) Stacked Ensemble Model

The stacked ensemble (stacking) approach was employed to improve overall performance by integrating predictions from multiple base models.^66^ In this framework, outputs from individual baseline models are combined, leveraging complementary features to enhance classification accuracy. By aggregating predictions from diverse models, stacking improves both the generalizability and stability of the final model.

In this study, the stacked ensemble was constructed using RF, XGBoost, AdaBoost, Logit Boost, Decision Tree, SVM and KNN models as base learners. A multilayer perceptron (MLP) with two hidden layers of 75 neurons each was used as the meta-learner to integrate the outputs from the base classifiers and generate the final predictions.

### 4.4 Performance evaluation of ML models on South Asian and Western datasets

The performance metrics for South Asian, western datasets were assessed using 7 different classifiers (Random forest, Logitboost, Adaboost, XGboost, Decision tree, SVM, and KNN). The CRC-associated taxa abundance data were analyzed, and models were evaluated using 20-repeated 10-fold stratified cross-validation. For each ML model accuracy, sensitivity, F1-measure, precision, and AUC are calculated and used as the performance metrics (Table-S1).

The performance metrics of various ML classifiers across South Asian cohorts follows the order LGBM>RF>XGBoost > AdaBoost > LogitBoost > Decision Tree. The highest AUC value across all the classifiers is highlighted in bold. The performance metrics of various ML classifiers across western (USA) cohorts follows the order LGBM>RF>XGBoost > AdaBoost > LogitBoost > Decision Tree. The highest AUC value across all the classifiers is highlighted in bold

The ROC curve corresponding to the model with the highest AUC is presented in Figure 4. The ROC curves for the remaining ML models are presented in supplementary information (Figure S5).

**Figure 4:**
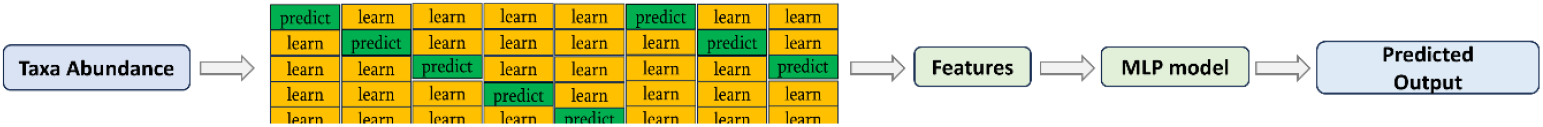
Conceptual diagram of the stacking ensemble framework, where outputs from diverse base learners are combined through a meta-model to enhance predictive accuracy.

**Table-1:** Performance evaluation of the various ML models on South Asian dataset.

| Model | Accuracy | Precision | Sensitivity | F1-Score | AUC | Specificity |
| --- | --- | --- | --- | --- | --- | --- |
| Random Forest | 0.744 ± 0.068 | 0.805 ± 0.123 | 0.547 ± 0.117 | 0.643 ± 0.101 | 0.813 ± 0.07 | 0.89 |
| Decision Tree | 0.703 ± 0.143 | 0.723 ± 0.143 | 0.535 ± 0.140 | 0.602 ± 0.115 | 0.700 ± 0.081 | 0.829 |
| AdaBoost | 0.734 ± 0.068 | 0.775 ± 0.125 | 0.560 ± 0.133 | 0.638 ± 0.104 | 0.795 ± 0.081 | 0.865 |
| LogitBoost | 0.739 ± 0.067 | 0.76 ± 0.119 | 0.571 ± 0.138 | 0.571 ± 0.138 | 0.798 ± 0.081 | 0.865 |
| XG Boost | 0.735±0.075 | 0.762 ± 0.121 | 0.580 ± 0.135 | 0.648 ± 0.108 | 0.812 ± 0.080 | 0.857 |
| SVM | 0.550 ± 0.077 | 0.487 ± 0.063 | 0.776 ± 0.147 | 0.593 ± 0.081 | 0.632 ± 0.116 | 0.380 |
| LGBM | 0.749 ± 0.070 | 0.773 ± 0.113 | 0.604 ± 0.121 | 0.670 ± 0.100 | **0.822 ± 0.067** | 0.859 |
| KNN | 0.578 ± 0.093 | 0.511 ± 0.117 | 0.531 ± 0.149 | 0.515 ± 0.119 | 0.605 ± 0.108 | 0.613 |

**Table-2:** Performance evaluation of the various ML models on western dataset.

| Model | Accuracy | Precision | Senstivity | F1-Score | AUC | Specificity |
| --- | --- | --- | --- | --- | --- | --- |
| Random Forest | 0.676 ± 0.042 | 0.675 ± 0.071 | 0.492 ± 0.101 | 0.563 ± 0.083 | **0.727 ± 0.051** | 0.817 |
| Decision Tree | 0.634 ± 0.070 | 0.616 ± 0.111 | 0.544 ± 0.102 | 0.524 ± 0.074 | 0.572 ± 0.066 | 0.602 |
| AdaBoost | 0.681 ± 0.063 | 0.730 ± 0.109 | 0.408 ± 0.140 | 0.514 ± 0.127 | 0.742 ± 0.054 | 0.888 |
| LogitBoost | 0.681 ± 0.034 | 0.709 ± 0.060 | 0.444 ± 0.071 | 0.544 ± 0.064 | 0.700± 0.061 | 0.860 |
| XGBoost | 0.679 ± 0.061 | 0.651±0.106 | 0.597±0.094 | 0.615±0.069 | 0.725± 0.057 | 0.741 |
| SVM | 0.667 | 0.681 | 0.402 | 0.499 | 0.67 | 0.86 |
| LGBM | 0.649 | 0.58 | 0.55 | 0.56 | 0.69 | 0.722 |
| KNN | 0.611 | 0.57 | 0.36 | 0.442 | 0.61 | 0.79 |

### 4.4 Explainability of the ML models

The interpretability of a machine learning model, specifically its ability to reveal the relationship between input features and output predictions, is paramount for gaining a profound understanding of how the model classifies data. This transparency is particularly crucial in sensitive domains like biomedical research, especially when predicting cancer from controls. In the context of cancer prediction, interpretability not only aids in validating model decisions but can also uncover critical biomarkers and support clinical hypotheses.

Feature importance methods are key techniques that quantify the individual contribution of each feature to the model’s prediction. For instance, the global feature importance derived from ensemble models like XGBoost are shown in Figure 5. It provides valuable insights into the most influential biological markers or patient characteristics considered by the model for distinguishing between CRC and controls. The top-ranked features for ML models considered (RF, DT, Adaboost, SVM, KNN, LGBM) are provided in the supplementary information (Supplementary Figure-S7-S20). Notably, *Peptostreptococcus, Parvimonas, Faecalibacterium*, and *Blautia* consistently emerged among the top 20 features across the models analyzed. The Shap analysis for XGBoost model for South Asian and Western cohorts is shown in the Figure 5. Understanding these features are vital for translating predictive models into actionable knowledge, ultimately supporting early diagnosis, guiding personalized treatment strategies, and advancing clinical applications.

**Figure 5:**
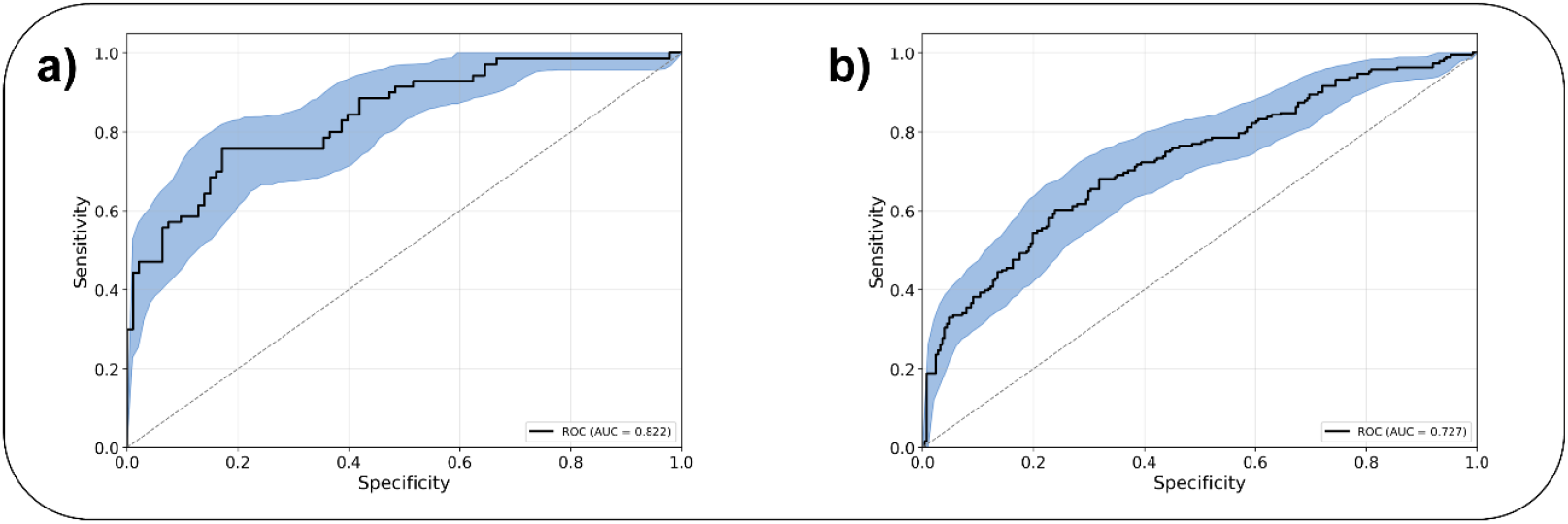
ROC Figure for the top performing ML models for the datasets considered a) South Asian b) Western.

The SHAP summary plots of feature importance derived from the XGBoost model for classifying CRC from controls across the South Asian and Western datasets are shown in Figure 6. It constructs an interpretable linear approximation of the model around each individual sample, enabling the quantification of feature contributions at the local level. By aggregating these local explanations across all samples, SHAP generates a global ranking of features that most strongly influence the model’s predictions. In these plots, the x-axis ranges from −1 to +1, with blue indicating negative contributions and red indicating positive contributions. A positive SHAP value for a feature (e.g., a bacterial genus) suggests a higher probability of CRC, whereas a negative SHAP value indicates a lower probability.

**Figure 6:**
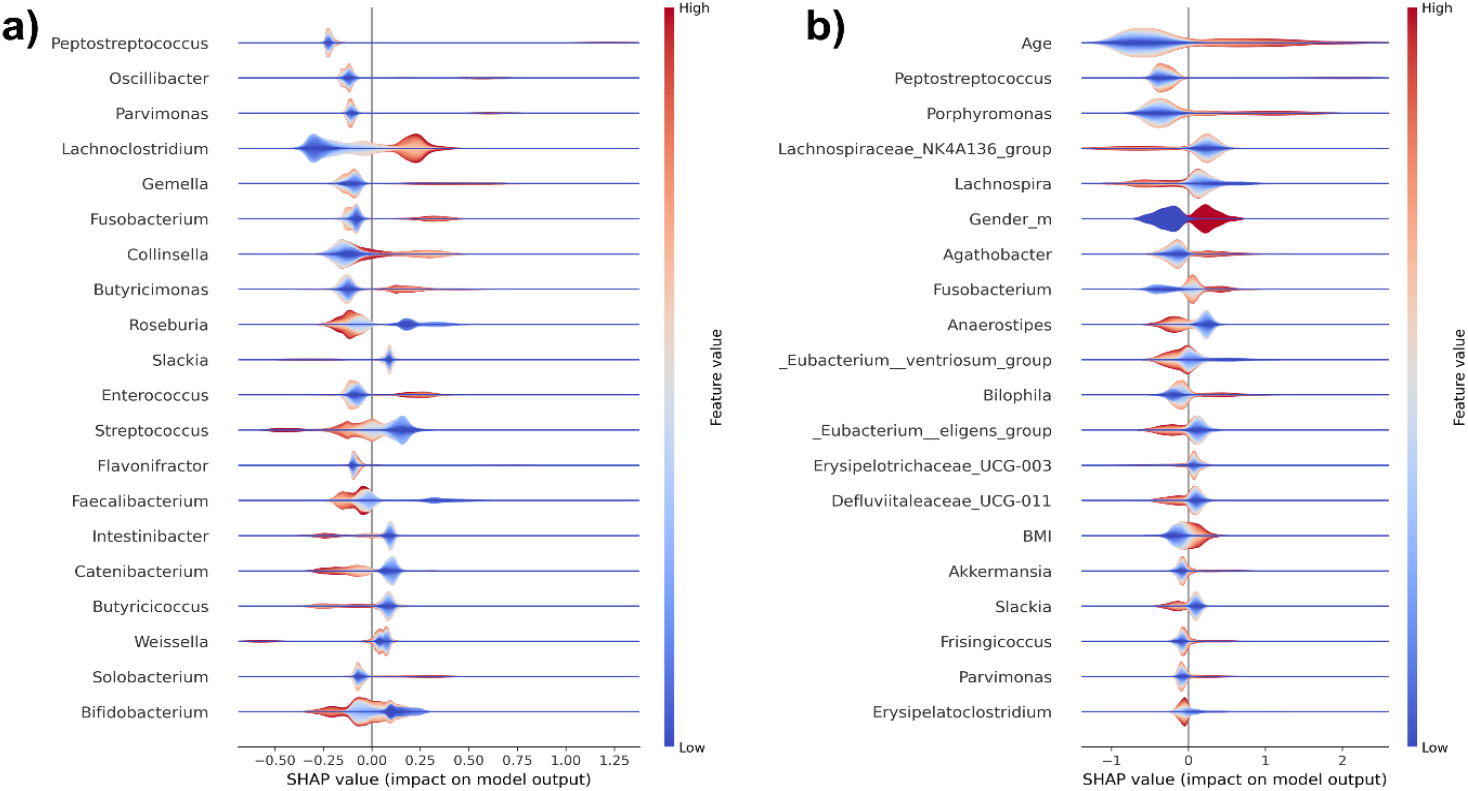
Shap summary plot illustrating the top 20 features ranked by their importance. Each point represents a subject’s Shapley value, with y-axis showing the feature names and the x-axis showing the corresponding Shapley values. The color gradient represents feature values (low to high), and features are ordered by their mean importance, with most influential features displayed at the top. A) South Asian dataset, B) Western dataset

**Figure 7:**
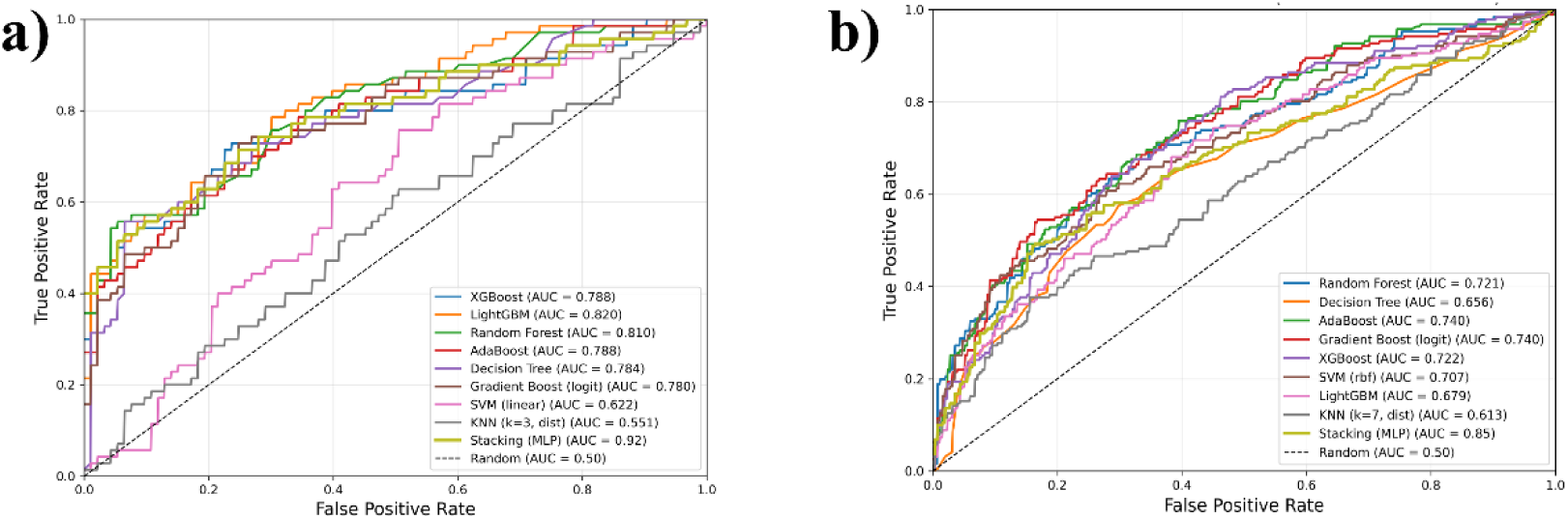
ROC curves of base models and stacked ensemble a) South Asian b) Western.

### 4.5 Stacking Ensemble Model

To enhance the predictive performance of our machine learning (ML) framework, we implemented a stacked ensemble learning approach that integrates multiple diverse algorithms. In this method, several base models are trained independently on features obtained from the base model. The predictions from these base models are then used as input features for a meta-learner, which combines them to generate the final prediction. This approach enables the ensemble to capture complex relationships that may not be fully addressed by individual models, thereby improving overall accuracy. In this study, we employed Random Forest, Decision Tree, AdaBoost, LogitBoost, and XGBoost as base classifiers. As the contribution of each model can vary—where some models may positively or negatively influence the final output—various combinations and configurations were explored to optimize the ensemble’s performance.

## 5. Discussion

In this study, we compared South Asian cohorts with Western cohorts to investigate microbiome variation in CRC and controls. Principal Coordinates Analysis (PCoA) revealed that geographical location exerted a stronger influence on gut microbiome composition than disease status. This finding underscores the need for more geographically diverse studies to fully elucidate the role of the microbiome in health and disease. Across cohorts, *Prevotella* was the predominant genus in South Asian populations, whereas *Bacteroides* dominated in Western cohorts. In addition, *Bifidobacterium* and *Faecalibacterium* were more abundant in controls, suggesting that their presence may be associated with a reduced risk of developing CRC.

Across all models evaluated, machine learning approaches successfully distinguished CRC cases from controls, albeit with varying accuracy. Consistent with previous studies, the Random Forest (RF) classifier achieved the highest overall performance across datasets, underscoring its robustness in analyzing complex, high-dimensional microbiome data. Although LGBM marginally outperformed RF in the South Asian dataset, the difference was negligible. The superior performance of RF likely stems from its ensemble framework, which combines predictions from multiple decision trees to reduce overfitting and model non-linear relationships between microbial features and disease status. Among all ML models tree based models (RF, DT, AdaBoost, XGBoost, LGBM) performed better for microbiome data than the SVM, KNN models which are distance based classifiers. Furthermore, RF’s resilience to noise and ability to manage correlated variables make it particularly well-suited for microbiome datasets, where feature interdependence is a common characteristic.

In the South Asian cohort, *Peptostreptococcus* and *Porphyromonas* (red points) displayed higher relative abundance on the positive side of the x-axis in the SHAP analysis (Figure 5), whereas lower relative abundance (blue points) appeared on the negative side. This pattern indicates that increased levels of these taxa are generally associated with a higher probability of CRC, while reduced abundance corresponds to a lower probability which is in line with previously reported studies.^67,68^ In contrast, *Collinsella* demonstrated the reverse trend, where higher abundance was linked to decreased CRC probability, as reflected on the negative side of the SHAP plot.

In the Western dataset, *Porphyromonas* and *Peptostreptococcus* were positioned on the positive side of the SHAP plot which is inline with earlier studies linking them to CRC as reported in earlier studies, while *Lachnospira* appeared on the negative side.^67,68^ Importantly, the top-ranked features identified by other ML models showed considerable overlap with those highlighted by SHAP analysis. Additionally, the SHAP plots emphasized key metadata variables—including age, gender, and BMI—among the top 20 influential features.

To improve model interpretability, SHAP-based top feature selection was applied to all classifiers, enabling the identification of microbial taxa most influential in driving predictions and offering insights into potential biomarkers for CRC detection. Among the top 20 features, *Fusobacterium, Porphyromonas, Peptostreptococcus*, and *Parvimonas* consistently emerged as potential microbiological markers, as indicated by both explainable AI (XAI) analysis and feature selection methods. The positive correlation of these taxa with model attribution toward the CRC class suggests their strong association with disease status. Notably, *Fusobacterium* has been previously implicated in CRC through multiple mechanisms, including the induction of genetic and epigenetic abnormalities, promotion of microsatellite instability, alteration of the immune microenvironment, and facilitation of metastasis, collectively contributing to tumorigenesis and disease progression. These findings reinforce its potential as a biomarker for identifying individuals at elevated risk of CRC.

Stacking ensemble methods combine the outputs of multiple base learning (ML) classifiers to leverage their complementary strengths and has improved overall predictive performance with MLP as metalearner. In our analysis, stacking enhanced the performance compared to based models, yielding better prediction with AUC greater than 90 % compared to their respective base models. This suggests that combining stacking ensemble with interpretable analysis could be used to build a generalizable model to predict CRC status.

## 6. Conclusion

This study underscores the importance of conducting microbiome investigations across diverse geographical regions to capture population-specific microbial signatures associated with CRC. By comparing South Asian and Western cohorts, we highlight how microbial diversity varies across demographics, emphasizing the need for population-specific studies. Machine learning classifiers successfully identified key bacterial taxa that differentiate CRC patients from controls, while explainable AI (XAI) analysis combined with a stacked ensemble approach further improved classification performance over individual base learners. These findings support the identification of robust and generalizable microbial biomarkers for early CRC detection. Furthermore, extending these methods to species-level analyses in future research may provide greater resolution for distinguishing CRC from controls, ultimately contributing to the development of reliable diagnostic tools.

## Acknowledgements

The authors would like to acknowledge International CRC Microbiome Network (AMS/CRUK) and Academy of Medical Sciences Global Challenges Research Fund Networking Grant (GCRFNG\100433); DBT/Wellcome Trust India alliance-Team Science Grants for funding (PEACOCC; Grant number: IA/TSG/23/1/600489).

## CRediT author statement

MMS, MB & AG: Conceptualization, Methodology; Software MMS, AG., VK; Data curation, Writing-Original draft preparation. MM, MB, NMS, AG, VK: Visualization, Investigation. MMS, AG; Supervision.: MB, NMS, RAS; Writing-Reviewing and Editing: MB, NMS, MMS.

## Consent for publication

Not applicable.

