## Supplementary file for "Integrative Microbiome Profiling of Colorectal Cancer Across South Asian and Western Cohorts Using Interpretable Machine Learning"

### MMS and AG equal first author contribution

#### Supplementary Information

#### Figure-S1

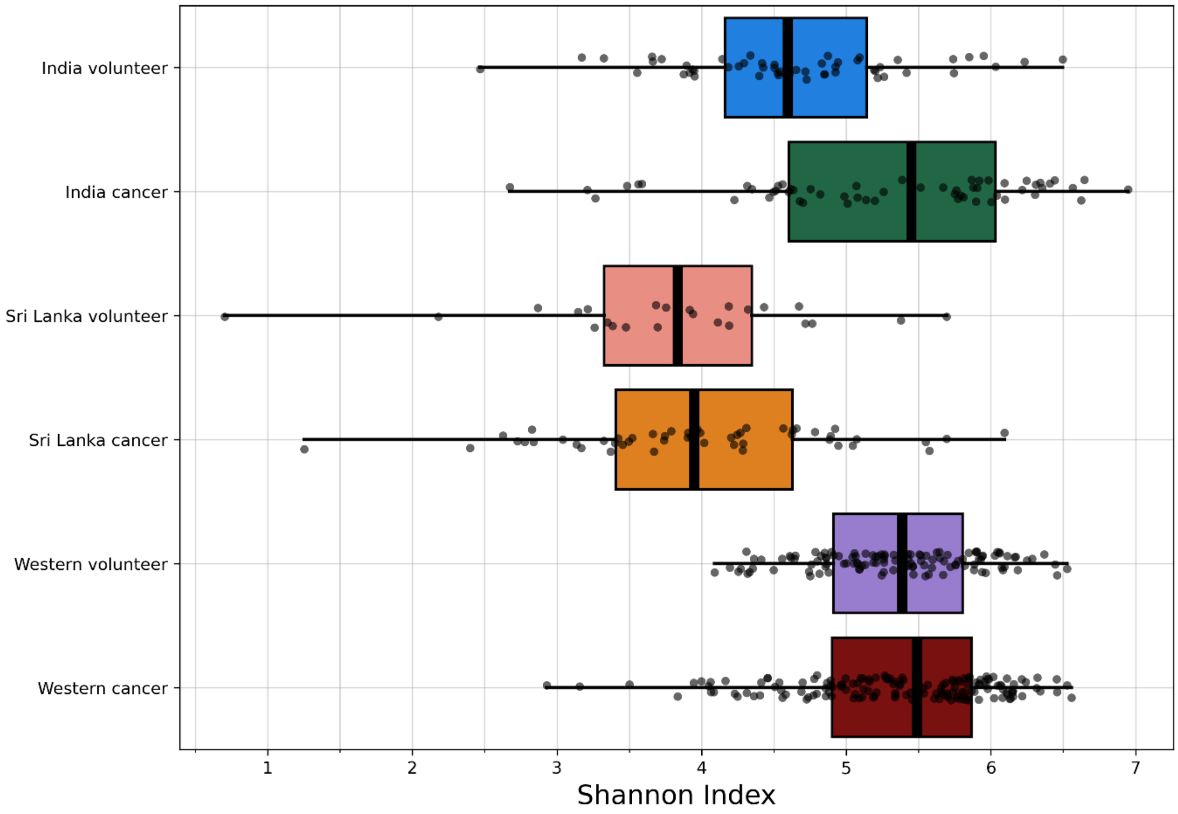


**Figure S1:** Shannon index alpha diversity for Indian, Sri Lankan and Western cohorts

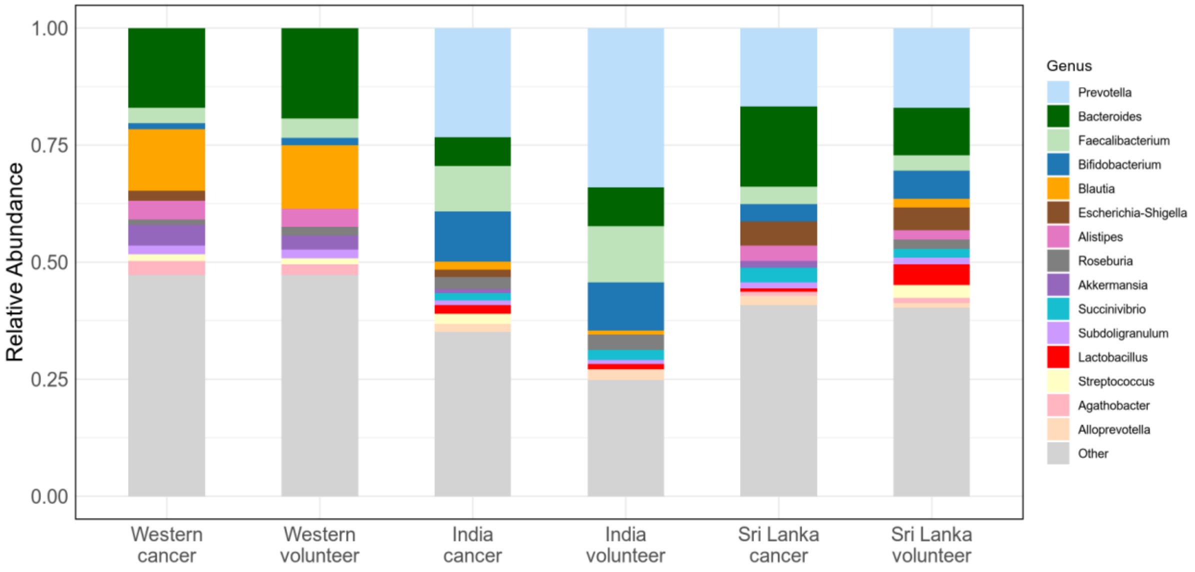


**Figure S2:** Taxa abundance showing top 15 genera across all the datasets considered

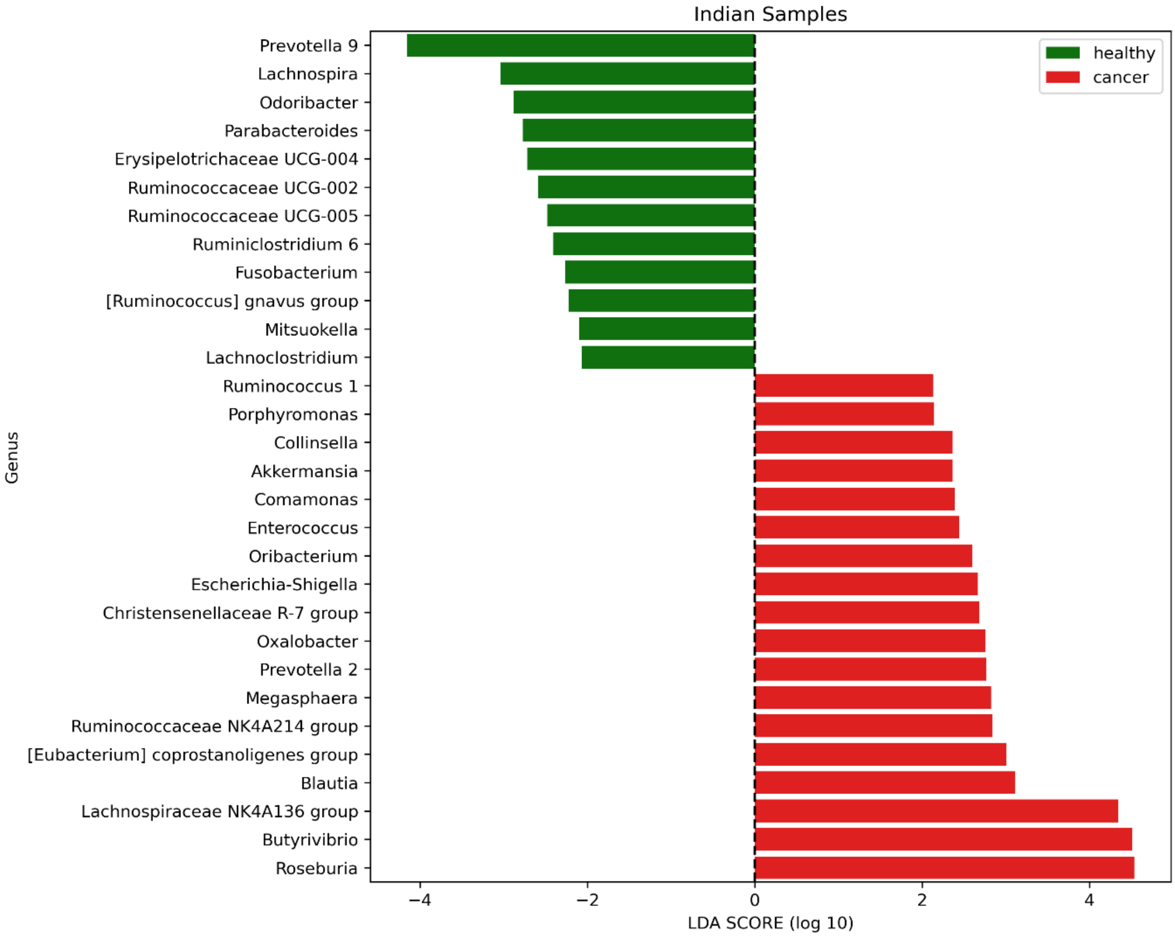


**Figure S3:** LefSe analysis of Indian Samples

#### Sri Lanka Samples

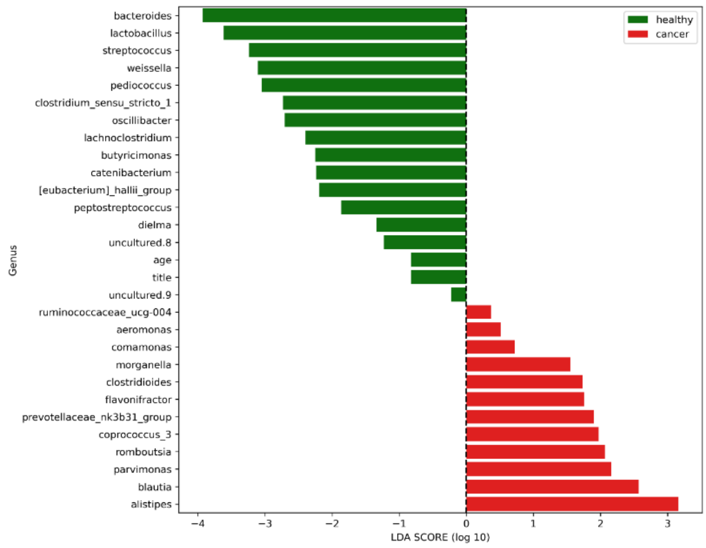


**Figure S4:** LefSe analysis of Sri Lankan Samples

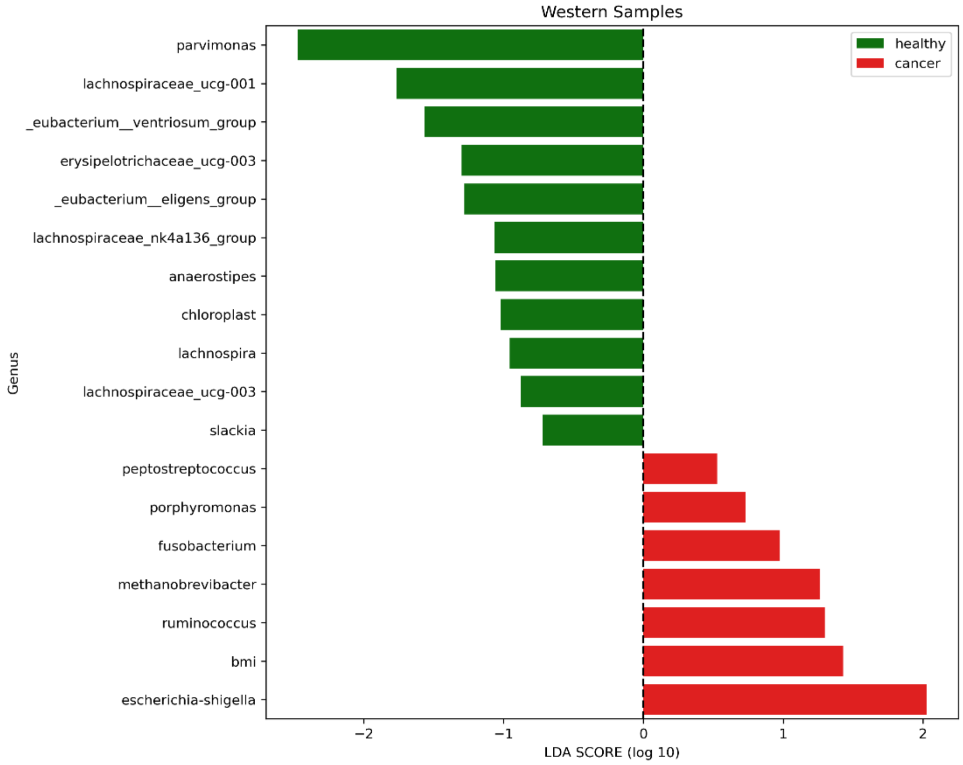


**Figure S4:** LefSe analysis of Western (USA) Samples

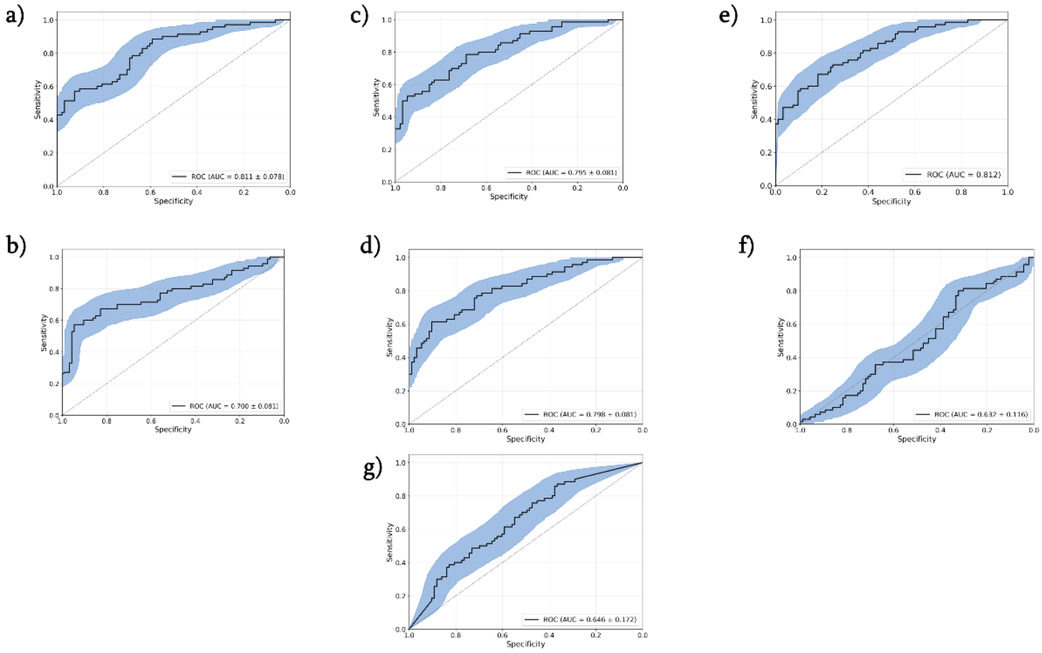


Figure S5: ROC curves for South Asia dataset a) Random forest, b) DT, c) Adaboost, d) LogitBoost e) XGBoost, f) SVM, g) KNN

**ROC curves for Western dataset :**

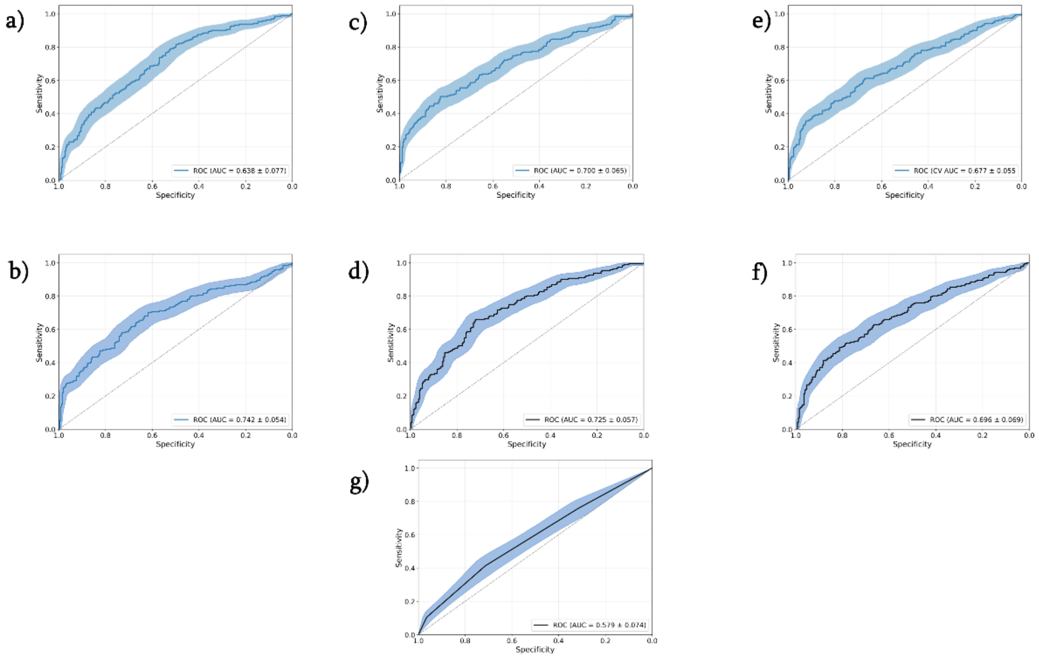

Figure S6: ROC curves for Western dataset  a) DT, b) Adaboost, c) LogitBoost d) XGBoost e) SVM f) LGBM, g) KNN

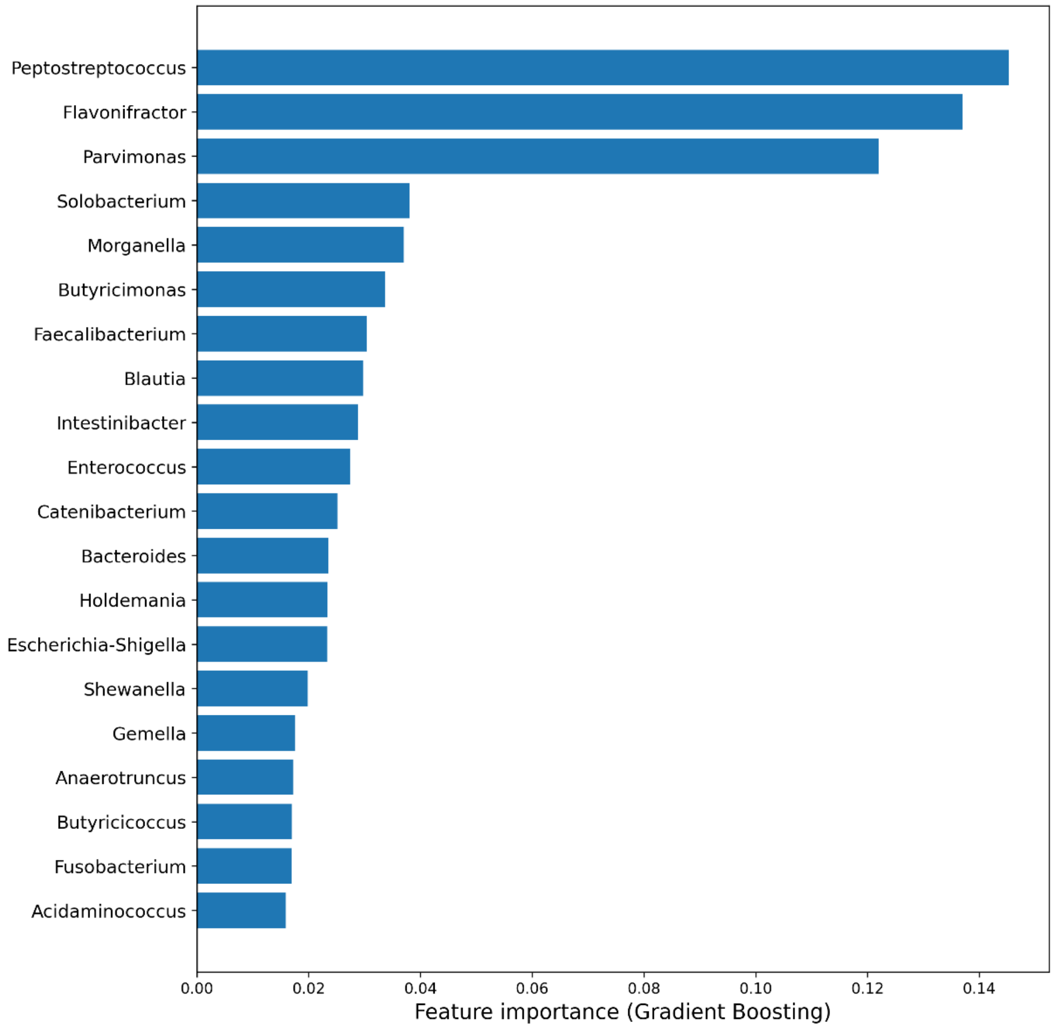


Figure S7:  Top-20 features extracted from logit boost ML model

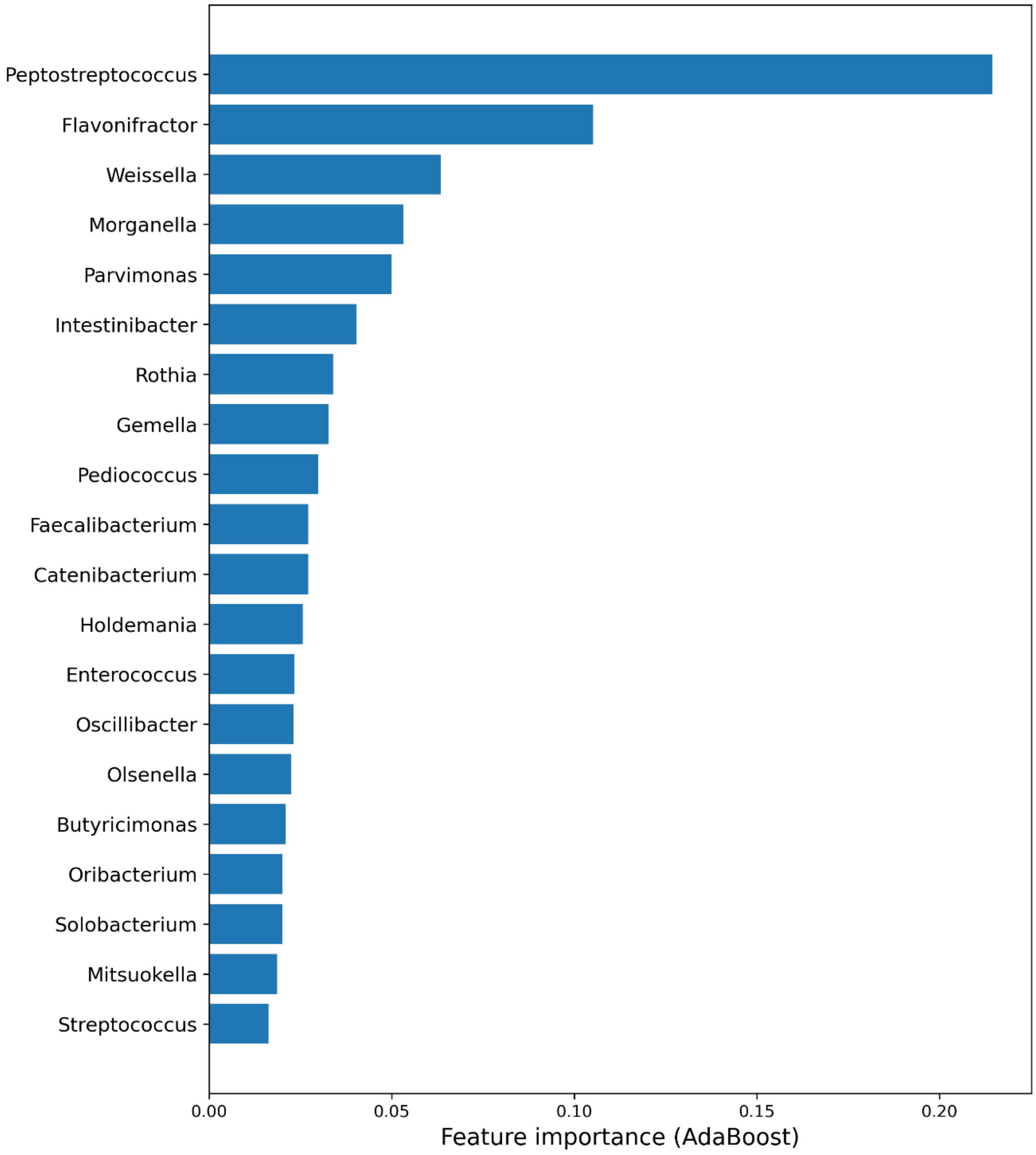


Figure S8: Top-20 features extracted from Ada boost ML model

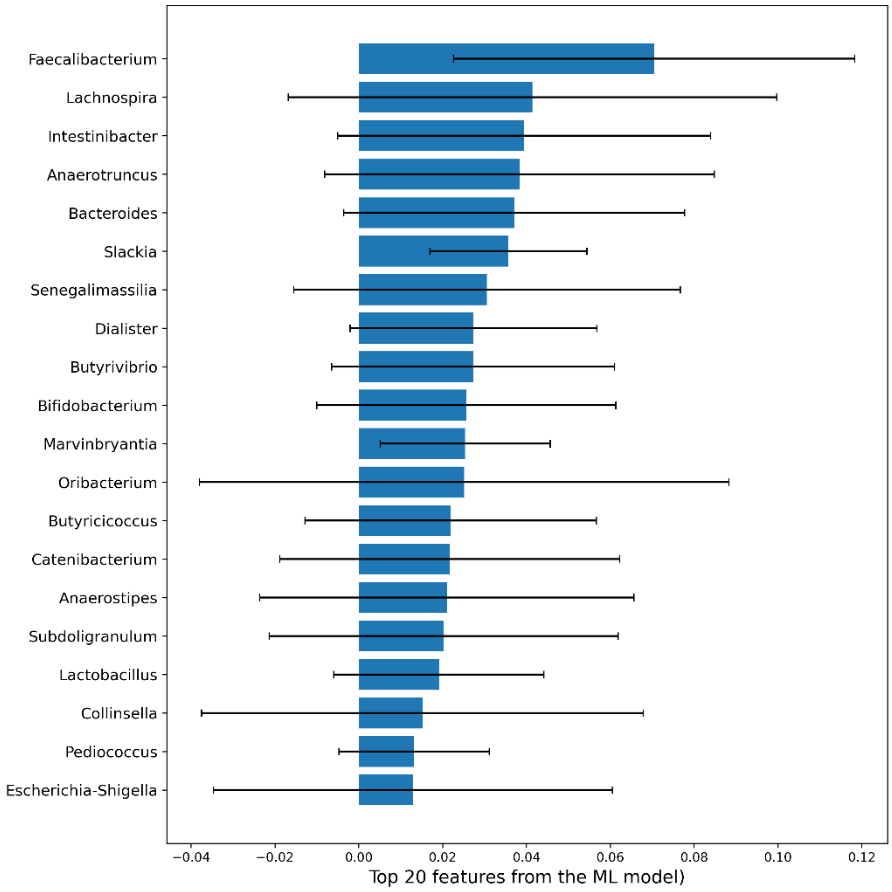


Figure S9: Top-20 features extracted from KNN ML model

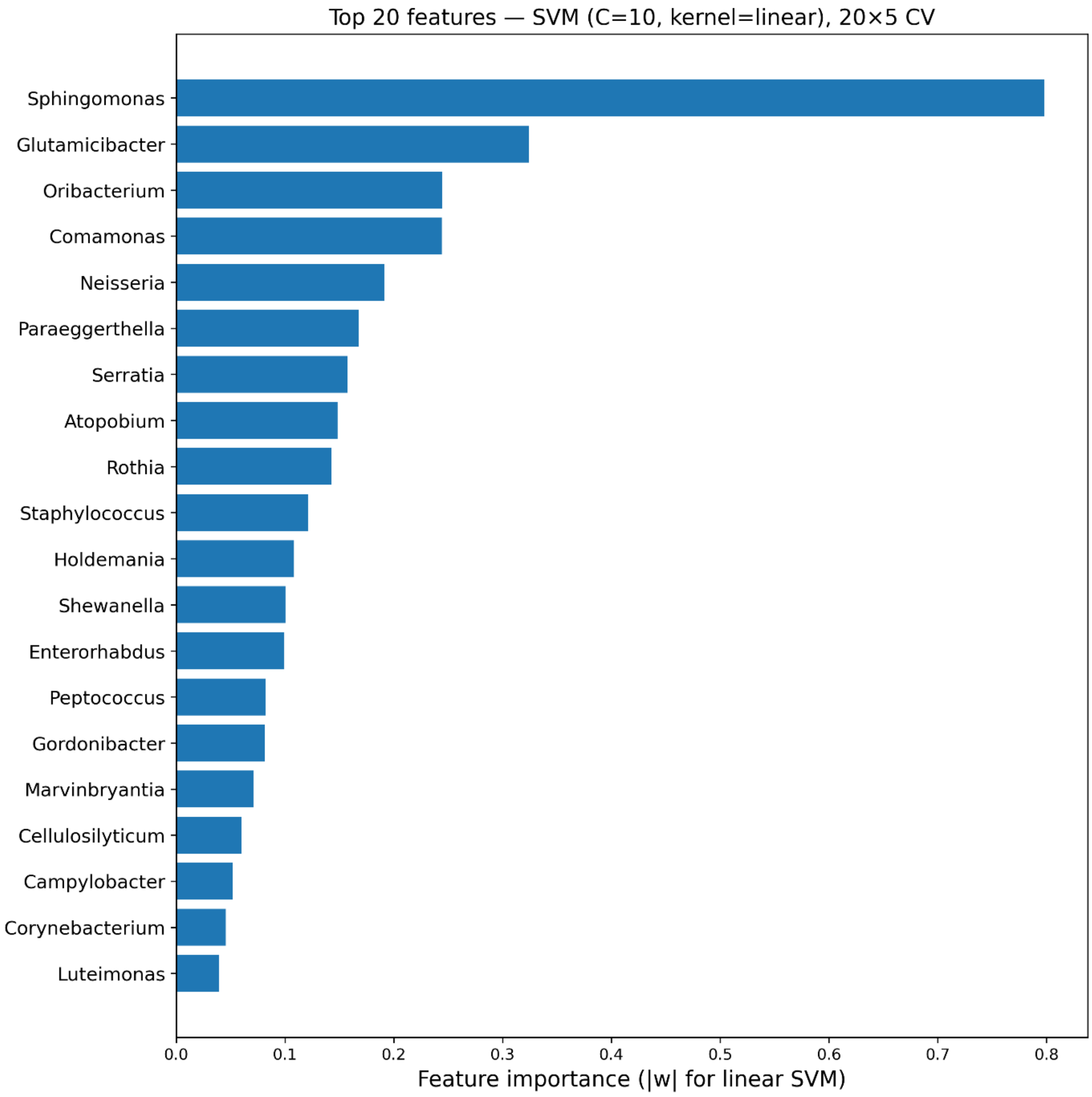


Figure S10: Top-20 features extracted from SVM ML model

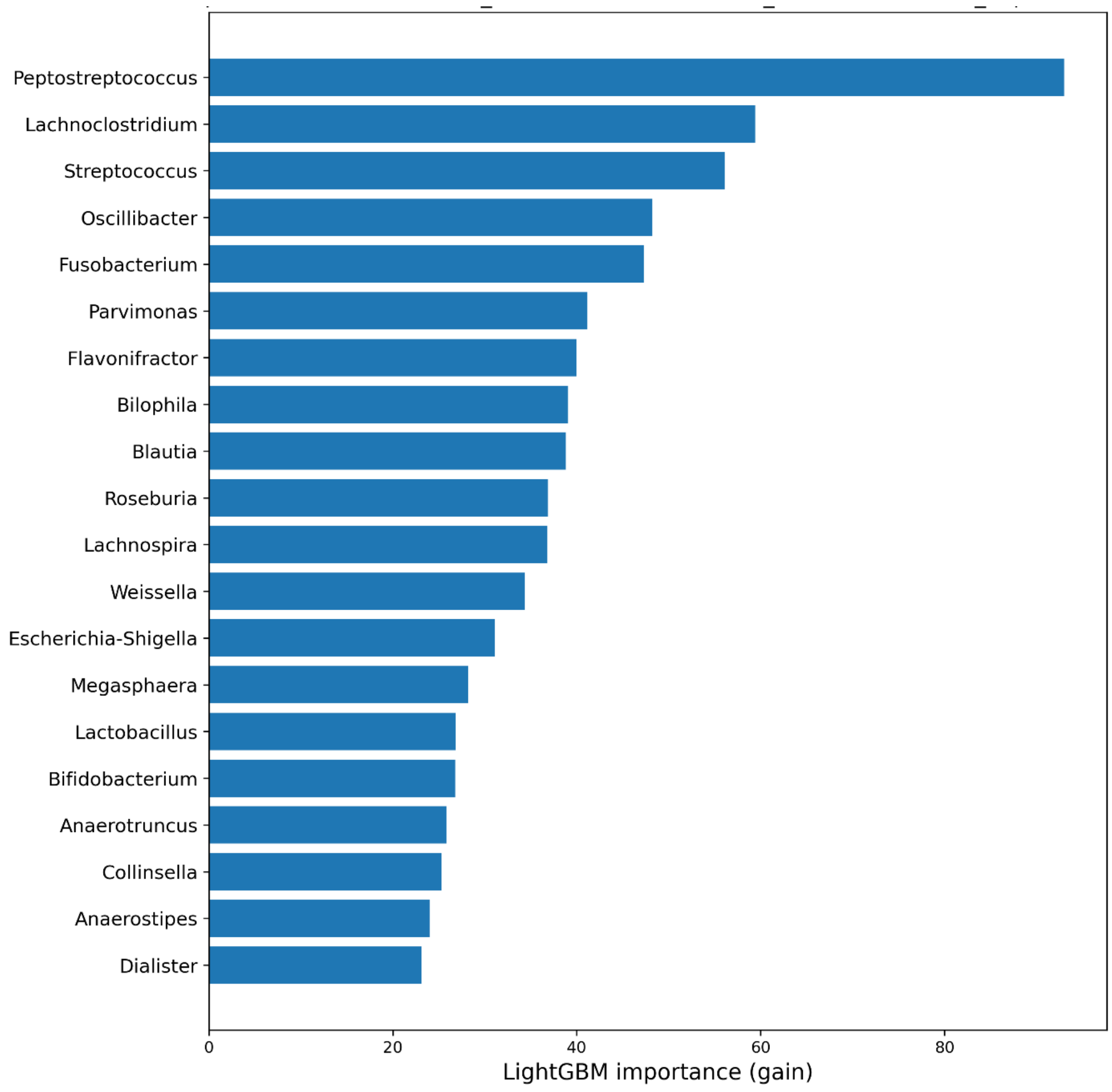


Figure S11: Top-20 features extracted from Light ML model

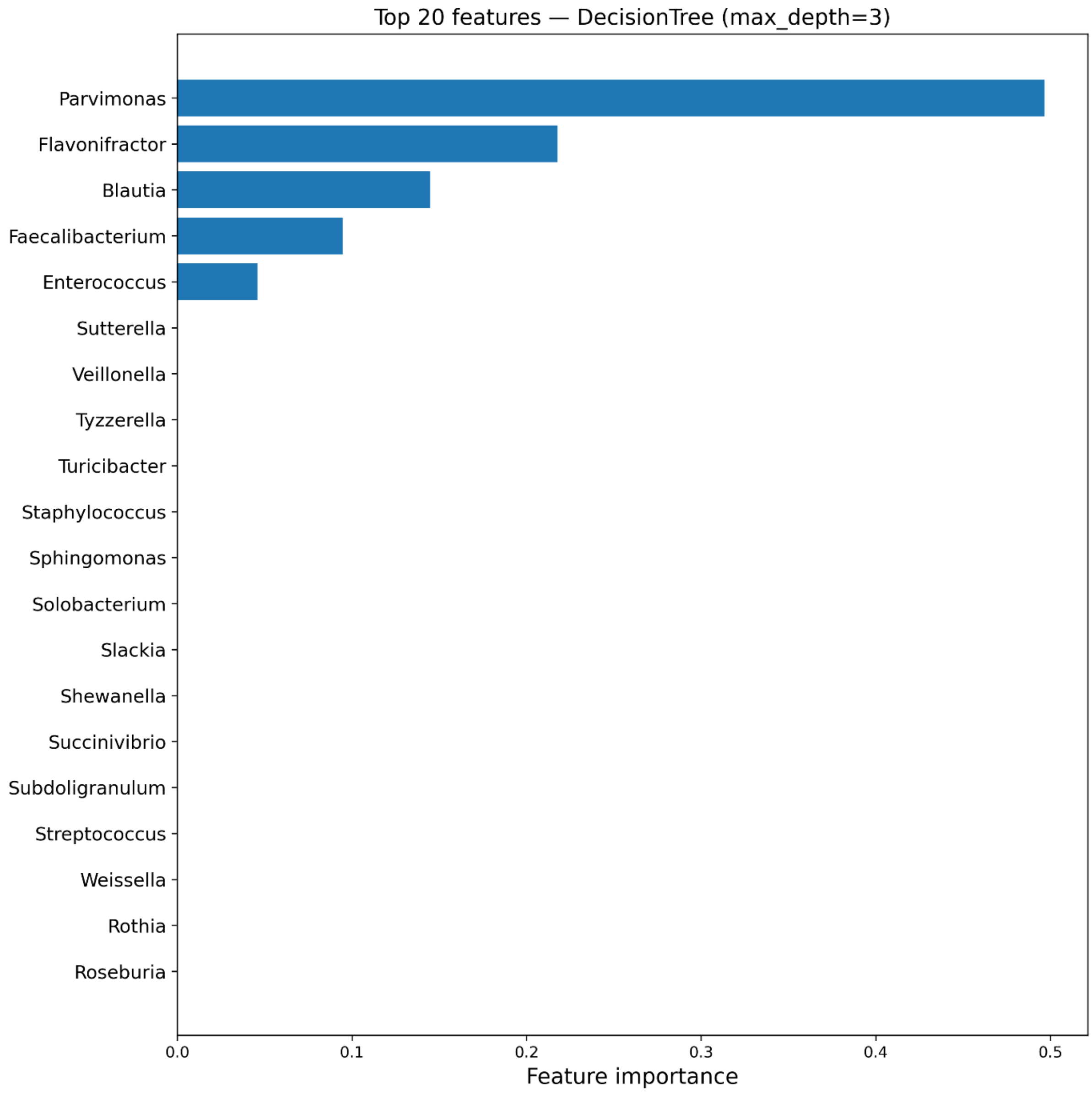


Figure S12: Top-20 features extracted from DT ML model

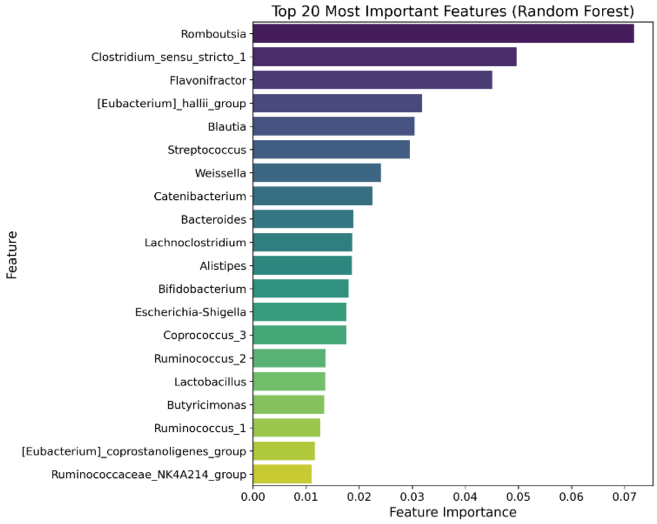


Figure S13: Top 20 features of considered by the RF model for distinguishing CRC from Non-CRC for Sri Lanka dataset

**Feature importance plot for Western data using different ML models:**

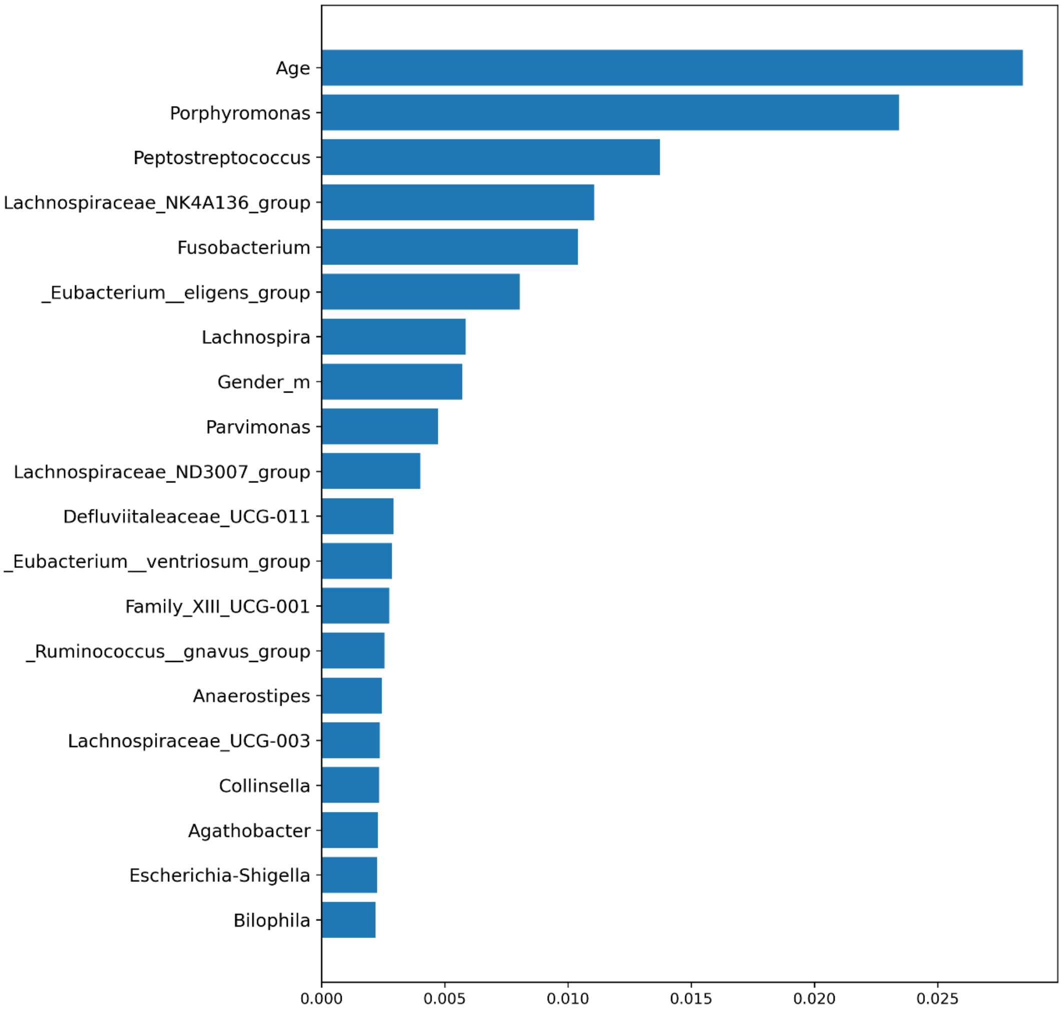


Figure S14: Top-20 features extracted from RF ML model

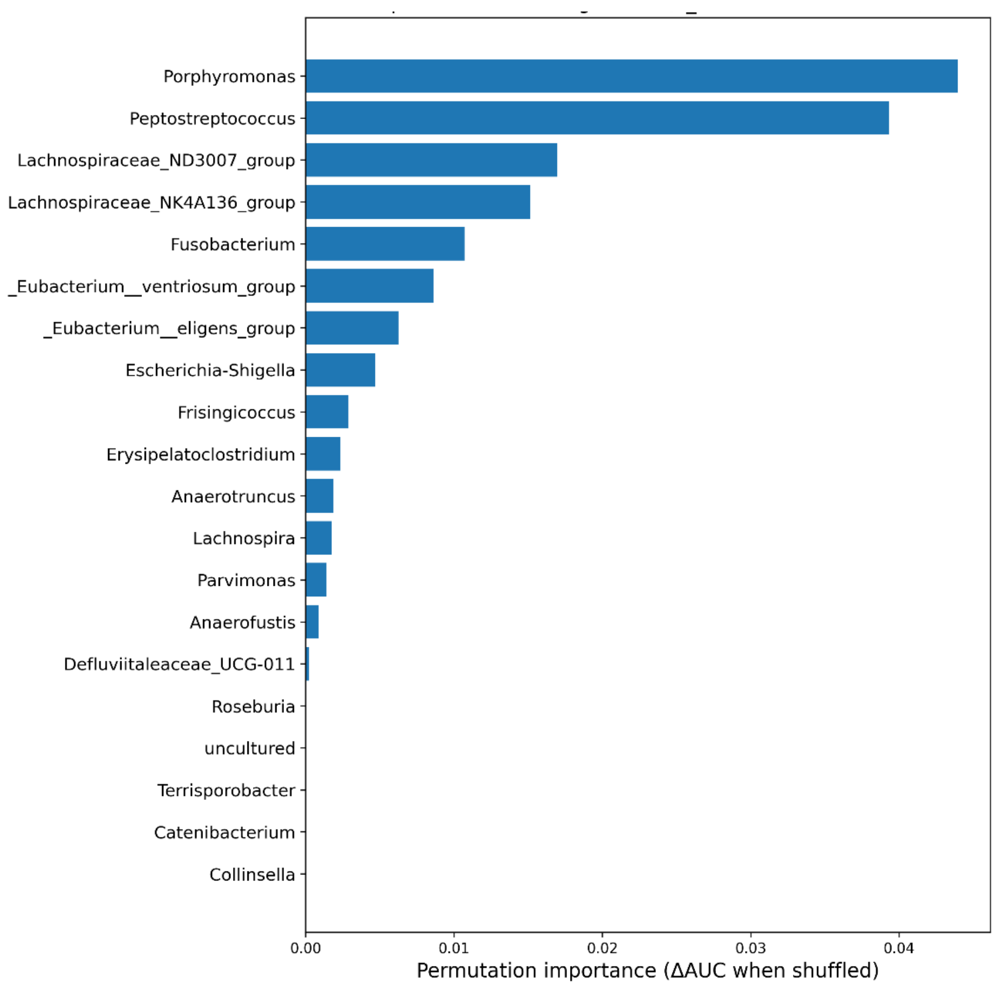


Figure S15: Top-20 features extracted from Logitboost ML model

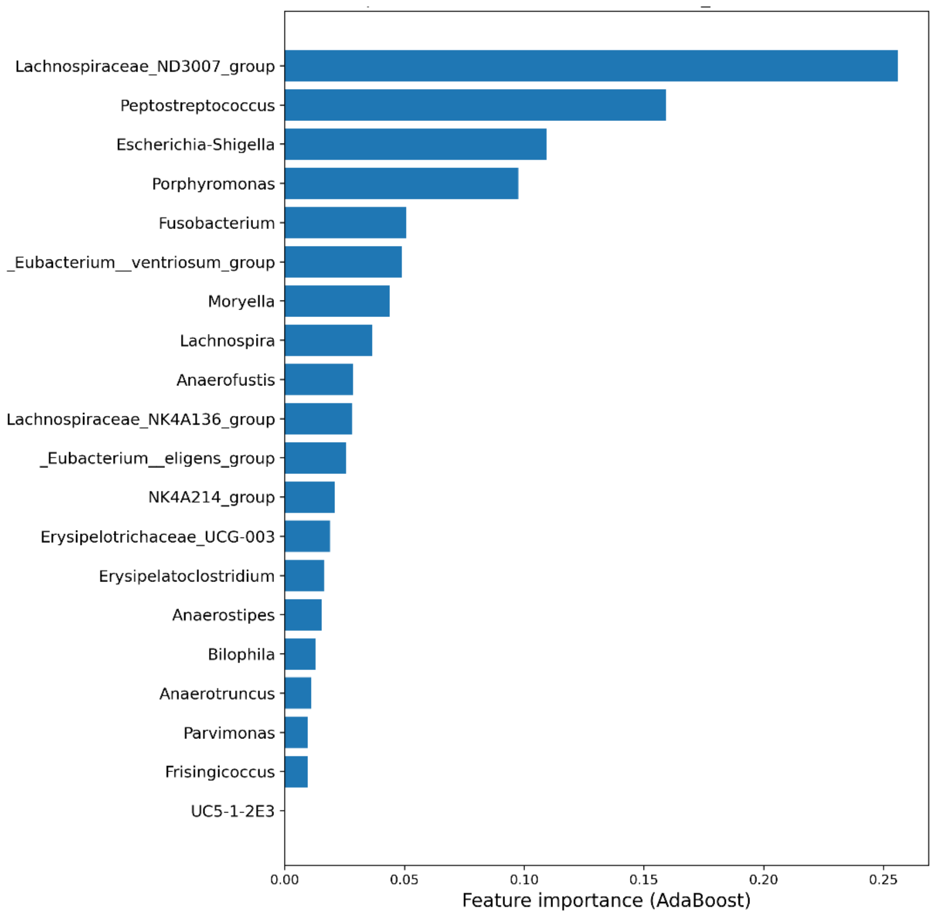


Figure S16: Top-20 features extracted from Adaboost ML model

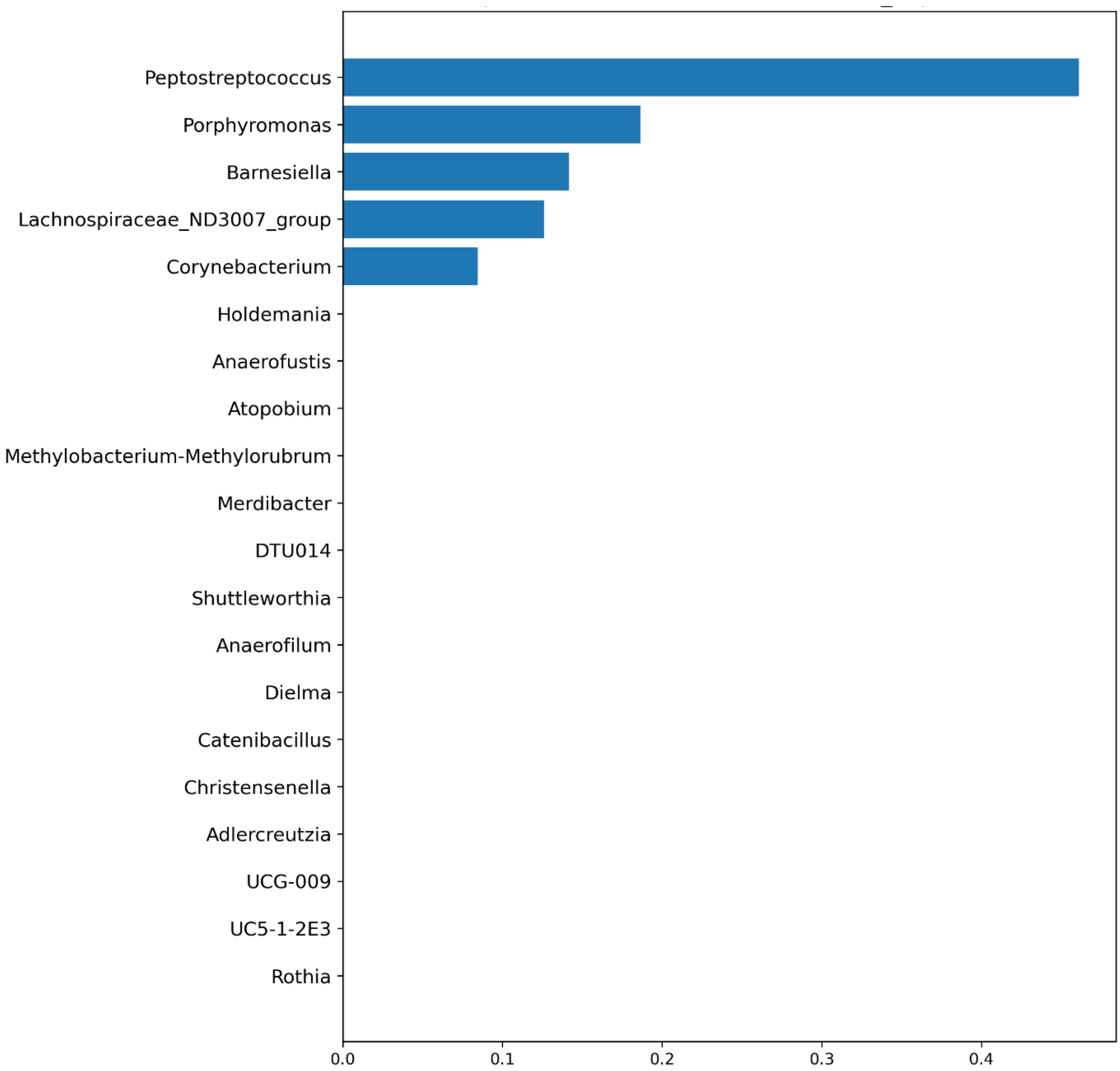


Figure S17: Top-20 features extracted from DT ML model

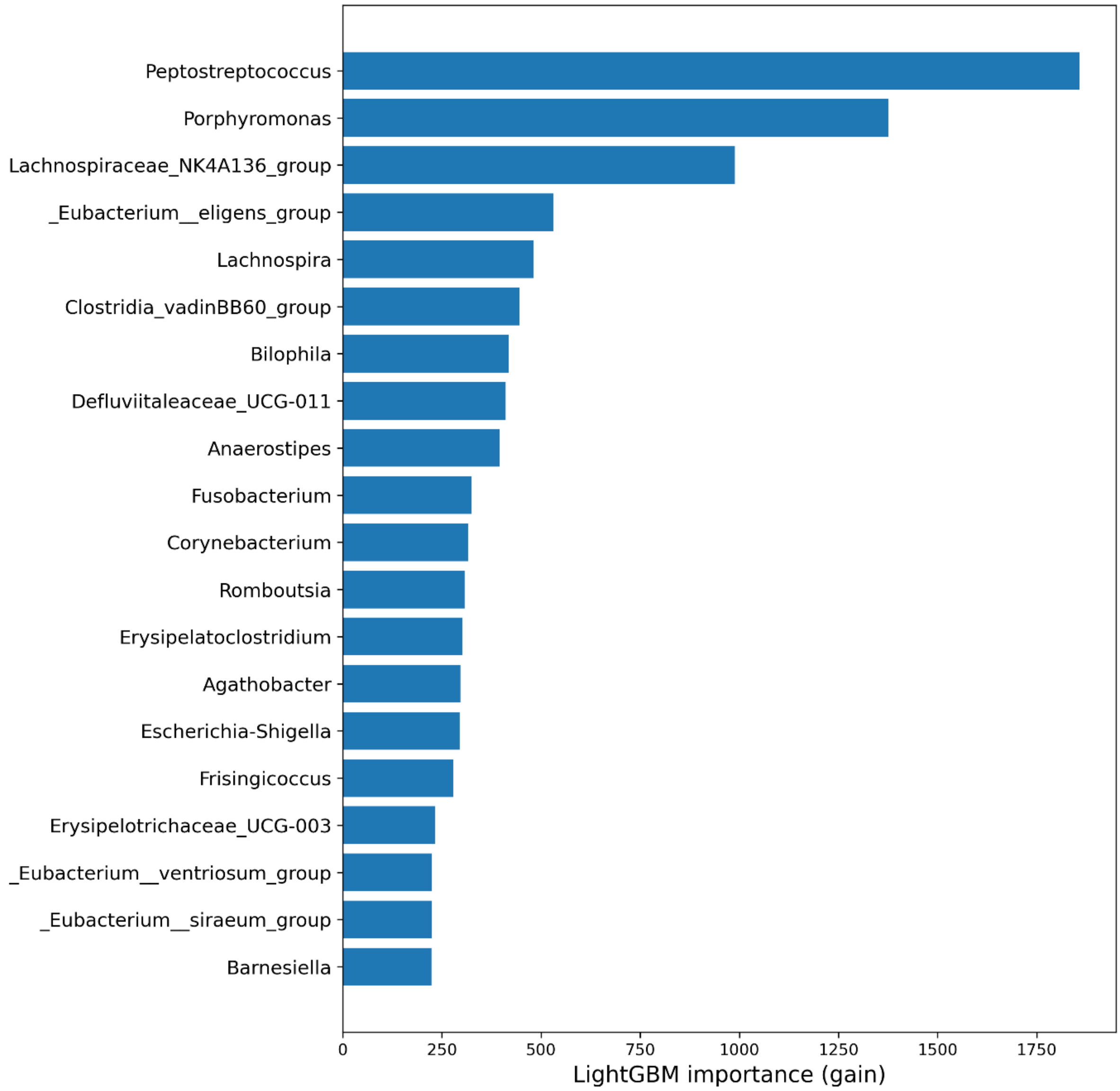


Figure S18: Top-20 features extracted from LightGBM ML model

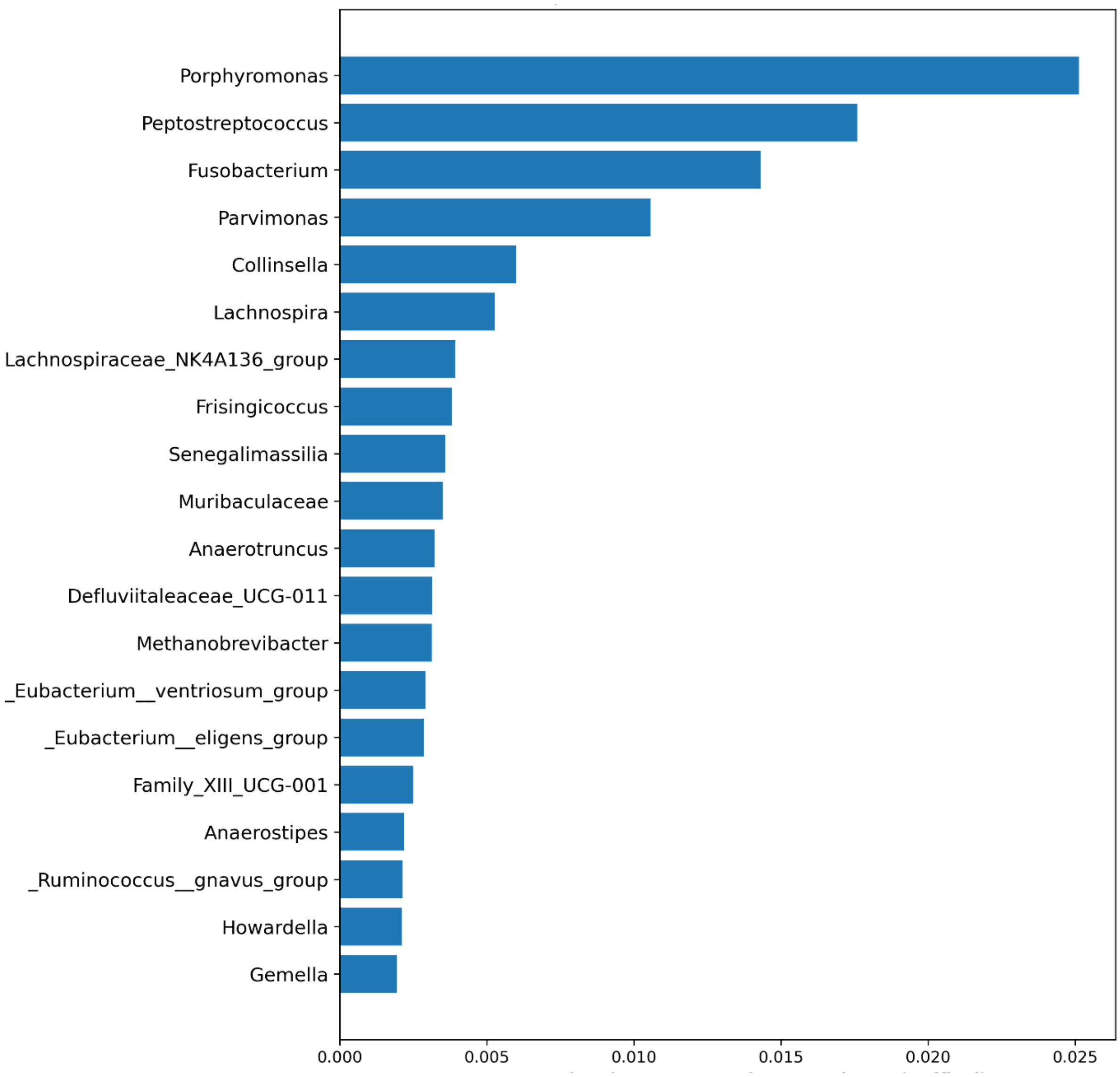


Figure S19: Top-20 features extracted from SVM ML model

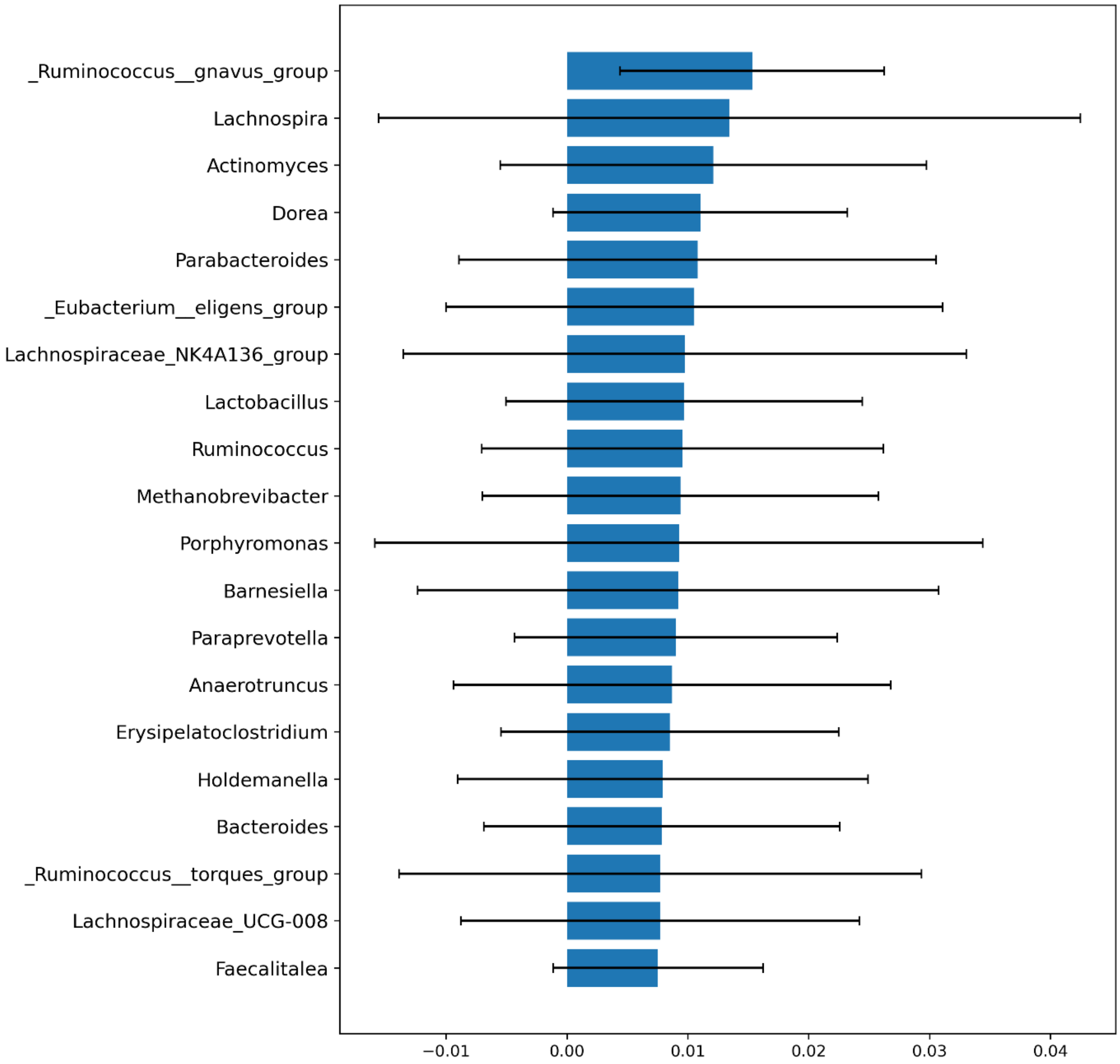


Figure S20: Top-20 features extracted from KNN ML model
